# Reaction times and other skewed distributions: problems with the mean and the median

**DOI:** 10.1101/383935

**Authors:** Guillaume A. Rousselet, Rand R. Wilcox

## Abstract

To summarise skewed (asymmetric) distributions, such as reaction times, typically the mean or the median are used as measures of central tendency. Using the mean might seem surprising, given that it provides a poor measure of central tendency for skewed distributions, whereas the median provides a better indication of the location of the bulk of the observations. However, the sample median is biased: with small sample sizes, it tends to overestimate the population median. This is not the case for the mean. Based on this observation, Miller (1988) concluded that “sample medians must not be used to compare reaction times across experimental conditions when there are unequal numbers of trials in the conditions.” Here we replicate and extend Miller (1988), and demonstrate that his conclusion was ill-advised for several reasons. First, the median’s bias can be corrected using a percentile bootstrap bias correction. Second, a careful examination of the sampling distributions reveals that the sample median is median unbiased, whereas the mean is median biased when dealing with skewed distributions. That is, on average the sample mean estimates the population mean, but typically this is not the case. In addition, simulations of false and true positives in various situations show that no method dominates. Crucially, neither the mean nor the median are sufficient or even necessary to compare skewed distributions. Different questions require different methods and it would be unwise to use the mean or the median in all situations. Better tools are available to get a deeper understanding of how distributions differ: we illustrate a powerful alternative that relies on quantile estimation. All the code and data to reproduce the figures and analyses in the article are available online.

## Introduction

Distributions of reaction times (RT) and many other quantities in social and life sciences are skewed (asymmetric) (Micceri, 1989; Limpert, Stahel & Abbt, 2001; Bono, Blanca, Arnau & Gómez-Benito, 2017). This asymmetry tends to differ among experimental conditions, such that a measure of central tendency and a measure of spread are insufficient to capture how conditions differ (Balota & Yap, 2011; Trafimow, Wang & Wang, 2018). Instead, to understand the potentially rich differences among distributions, it is advised to consider multiple quantiles of the distributions (K. Doksum, 1974; K. A. Doksum & Sievers, 1976; Pratte, Rouder, Morey & Feng, 2010; Rousselet, Pernet & Wilcox, 2017), or to explicitly model the shapes of the distributions (Heathcote, Popiel & Mewhort, 1991; Rouder, Lu, Speckman, Sun & Jiang, 2005; Palmer, Horowitz, Torralba & Wolfe, 2011; Matzke, Dolan, Logan, Brown & Wagenmakers, 2013). Yet, it is still common practice to summarise RT distributions using a single number, most often the mean: that one value for each participant and each condition can then be entered into a group ANOVA to make statistical inferences. Because of the skewness (asymmetry) of reaction times, the mean is however a poor measure of central tendency: skewness shifts the mean away from the bulk of the distribution, an effect that can be amplified by the presence of outliers or a thick right tail. For instance, in Figure 1A, the median better represents the typical observation than the mean because it is closer to the bulky part of the distribution. (Note that all figures in this article can be reproduced using code and data available in a reproducibility package on *figshare* (Rousselet & Wilcox, 2018a). Supplementary material notebooks referenced in the text are also available in this reproducibility package).

**Figure 1.**
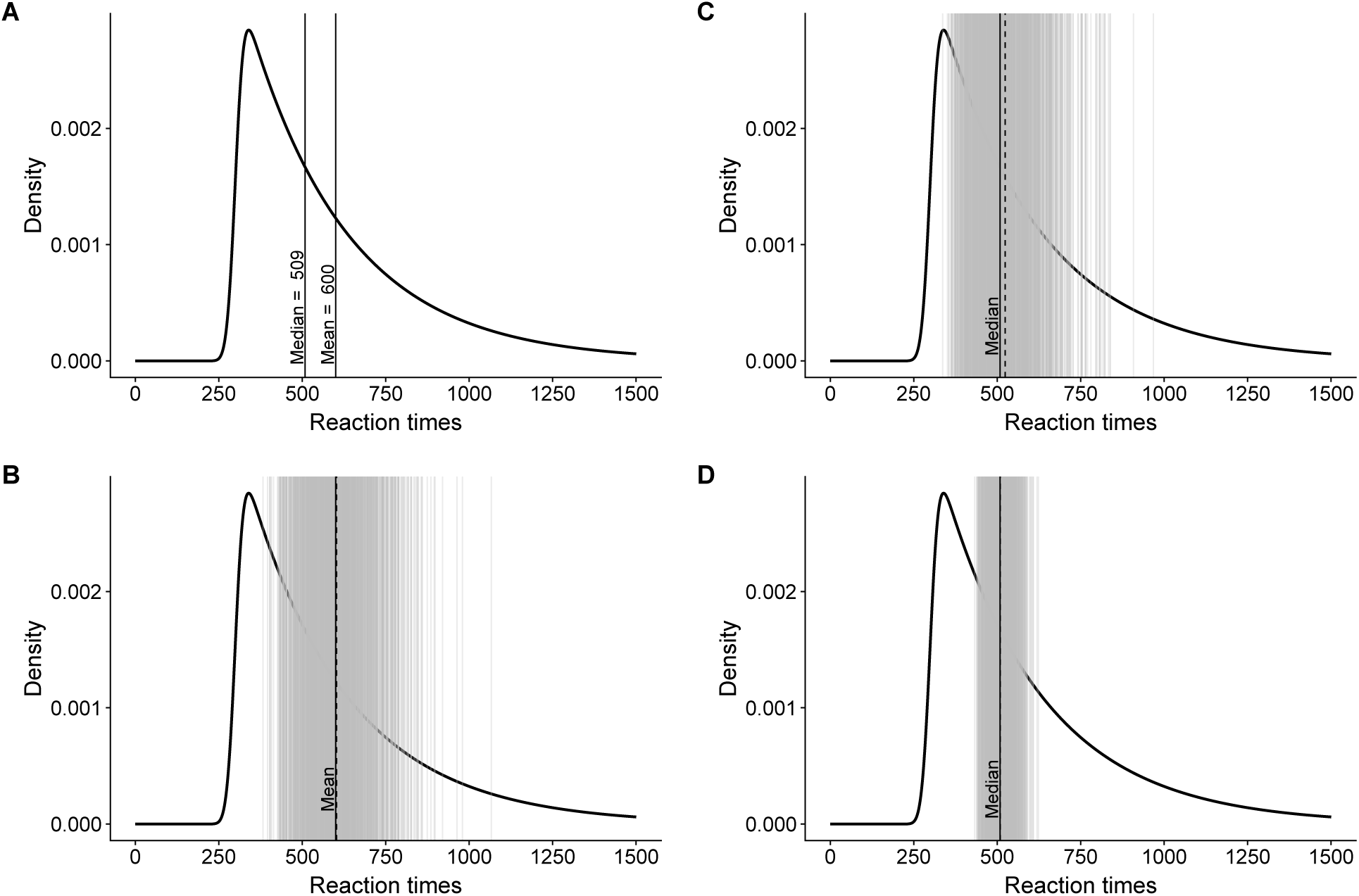
Skewness, sampling and bias. **A.** Ex-Gaussian distribution with parameters *µ* = 300,*σ* = 20 and *τ* = 300. The distribution is bounded to the left and has a long right tail. This distribution is used by convenience for illustration, because it looks like a very skewed reaction time distribution. The vertical lines mark the population mean and median. **B.** The vertical grey lines indicate 1,000 means from 1,000 random samples of 10 observations. As in panel A, the vertical black line marks the population mean. The vertical black dashed line marks the mean of the 1,000 sample means. **C.** Same as panel B, but for the median. **D.** Same as C, but for 1,000 samples of 100 observations.

So the median appears to be a better choice than the mean if the goal is to have a single value that reflects the location of most observations in a skewed distribution. In our experience the median is the most often used alternative to the mean in the presence of skewness. The choice between the mean and the median is however more complicated. Depending on the goals of the experimenter and the situation at hand, the mean or the median can be a better choice, but most likely neither is the best choice - no method dominates across all situations. For instance, it could be argued that because the mean is sensitive to skewness, outliers and the thickness of the right tail, it is better able to capture changes in the shapes of the distributions among conditions. As we will see, this intuition is correct in some situations. But the use of a single value to capture shape differences necessarily leads to intractable analyses because the same mean could correspond to various shapes. If the goal is to understand shape differences between conditions, a multiple quantile approach or explicit shape modelling should be used instead, as mentioned previously.

The mean and the median differ in another important aspect: for small sample sizes, the sample mean is unbiased, whereas the sample median is biased. Concretely, if we perform many times the same RT experiment, and for each experiment we compute the mean and the median, the average mean will be very close to the population mean. As the number of simulated experiments increases, the average sample mean will converge to the exact population mean. This is not the case for the median when sample size is small. However, over many studies, the median of the sample medians is equal to the population median. That is, the median is median unbiased (this definition of bias also has the advantage, over the standard definition using the mean, to be transformation invariant Efron and Hastie (2016)). In contrast, when dealing with skewed distributions, the sample mean is median biased.

To illustrate, imagine that we perform simulated experiments to try to estimate the mean and the median population values of the skewed distribution in Figure 1A. Let’s say we take 1,000 samples of 10 observations. For each experiment (sample), we compute the mean. These sample means are shown as grey vertical lines in Figure 1B. A lot of them fall very near the population mean (black vertical line), but some of them are way off. The mean of these estimates is shown with the black dashed vertical line. The difference between the mean of the mean estimates and the population value is called bias. Here bias is small (2.5). Increasing the number of experiments will eventually lead to a bias of zero. In other words, the sample mean is an unbiased estimator of the population mean.

For small sample sizes from skewed distributions, this is not the case for the median. If we proceed as we did for the mean, by taking 1,000 samples of 10 observations, the bias is 15.1: the average median across 1,000 experiments over-estimates the population median (Figure 1C). Increasing sample size to 100 reduces the bias to 0.7 and improves the precision of our estimates. On average, we get closer to the population median.

The reason for the bias of the median is explained by Miller (1988):

> ’Like all sample statistics, sample medians vary randomly (from sample to sample) around the true population median, with more random variation when the sample size is smaller. Because medians are determined by ranks rather than magnitudes of scores, the population percentiles of sample medians vary symmetrically around the desired value of 50%. For example, a sample median is just as likely to be the score at the 40th percentile in the population as the score at the 60th percentile. If the original distribution is positively skewed, this symmetry implies that the distribution of sample medians will also be positively skewed. Specifically, unusually large sample medians (e.g., 60th percentile) will be farther above the population median than unusually small sample medians (e.g., 40th percentile) will be below it. The average of all possible sample medians, then, will be larger than the true median, because sample medians less than the true value will not be small enough to balance out the sample medians greater than the true value. Naturally, the more the distribution is skewed, the greater will be the bias in the sample median.’

Because of this bias, Miller (1988) recommended to not use the median to study skewed distributions in certain situations. As we demonstrate here, the problem is more complicated and the choice between the mean and the median depends on the goal of the researcher. In this article, which is organised in 5 sections, we explore the advantages and disadvantages of the sample mean and the sample median. First, we replicate Miller’s simulations of estimations from single distributions. Second, we introduce bias correction and apply it to Miller’s simulations. Third, we examine sampling distributions in detail to reveal unexpected features of the sample mean and the sample median. Fourth, we extend Miller’s simulations to consider false and true positives for the comparisons of two conditions. Finally, we consider a large dataset of RT from a lexical decision task, which we use to contrast different approaches.

## Replication of Miller 1988

To illustrate the sample median’s bias, Miller (1988) employed 12 ex-Gaussian distributions that differed in skewness (Table 1). The distributions are illustrated in Figure 2, and colour-coded using the difference between the mean and the median as a non-parametric measure of skewness. Figure 1 used the most skewed distribution of the 12, with parameters (*µ* = 300, *σ* = 20, *τ* = 300).

**Table 1.**
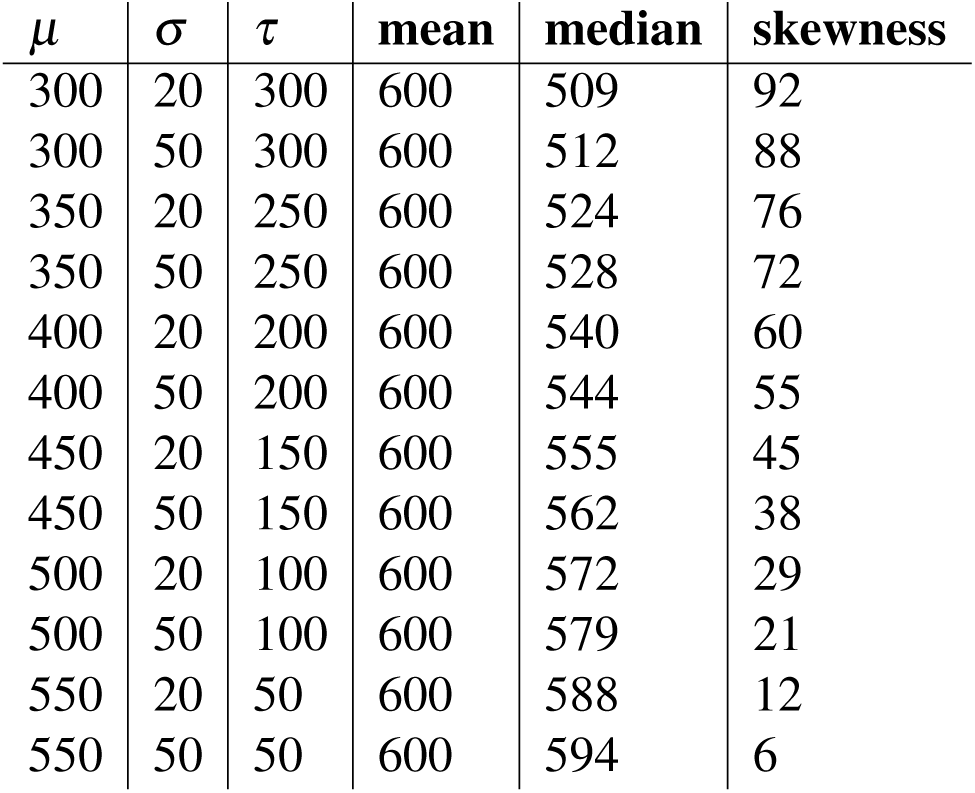
Miller’s 12 ex-Gaussian distributions. Each distribution is defined by the combination of the three parameters *µ* (mu), *σ* (sigma) and *τ* (tau). The mean is defined as the sum of parameters *µ* and *τ*. The median was calculated based on samples of 1,000,000 observations (Miller 1988 used 10,000 observations). Skewness is defined as the difference between the mean and the median.

**Figure 2.**
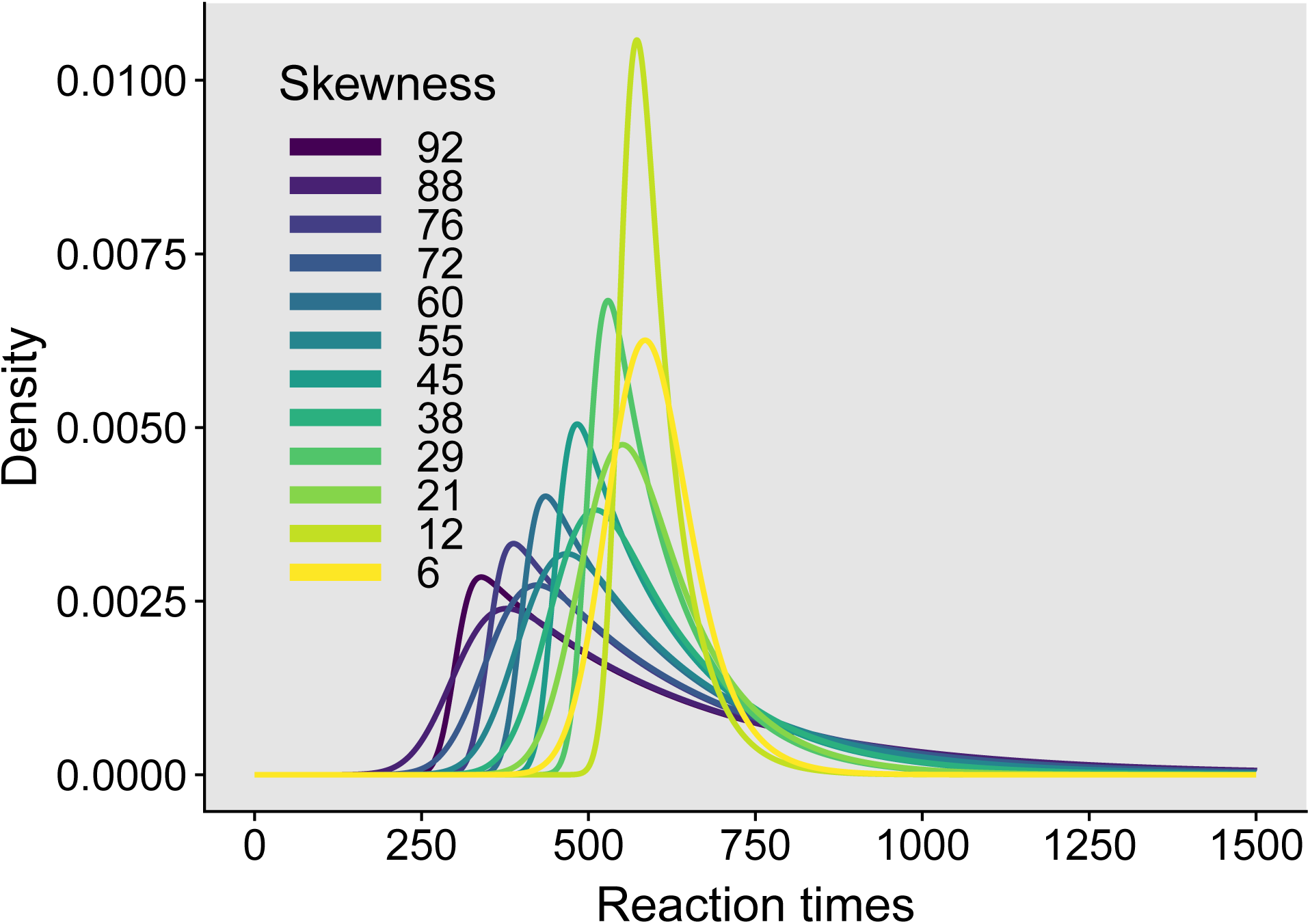
Miller’s 12 ex-Gaussian distributions.

To estimate bias, following Miller (1988) we performed a simulation in which we sampled with replacement 10,000 times from each of the 12 distributions. We took random samples of sizes 4, 6, 8, 10,15, 20, 25, 35, 50 and 100, as did Miller. For each random sample, we computed the mean and the median. For each sample size and ex-Gaussian parameter, the bias was then defined as the difference between the mean of the 10,000 sample estimates and the population value.

First, as shown in Figure 3, we can check that the mean is not biased. Each line shows the results for one type of ex-Gaussian distribution: the mean of 10,000 simulations for different sample sizes minus the population mean (600). Irrespective of skewness and sample size, bias is very near zero.

**Figure 3.**
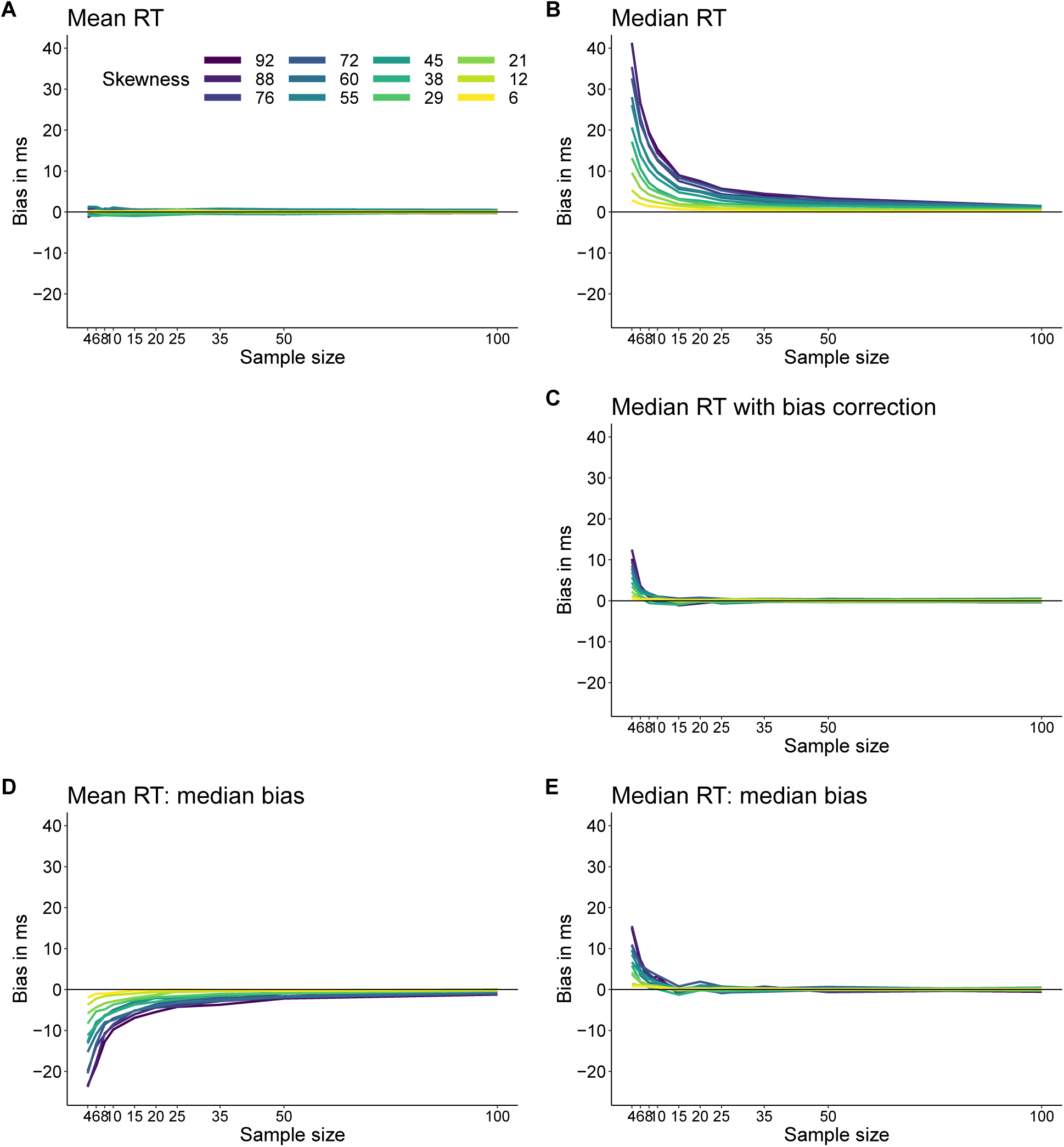
Bias estimation. **A.** Bias for mean reaction times. **B.** Bias for median reaction times. **C.** Bias for median reaction times after bootstrap bias correction. **D.** Median bias for mean reaction times. **E.** Median bias for median reaction times.

Contrary to the mean, the median estimates are biased for small sample sizes. The values from our simulations are very close to the values reported in Miller (1988) (Table 2).

**Table 2.**
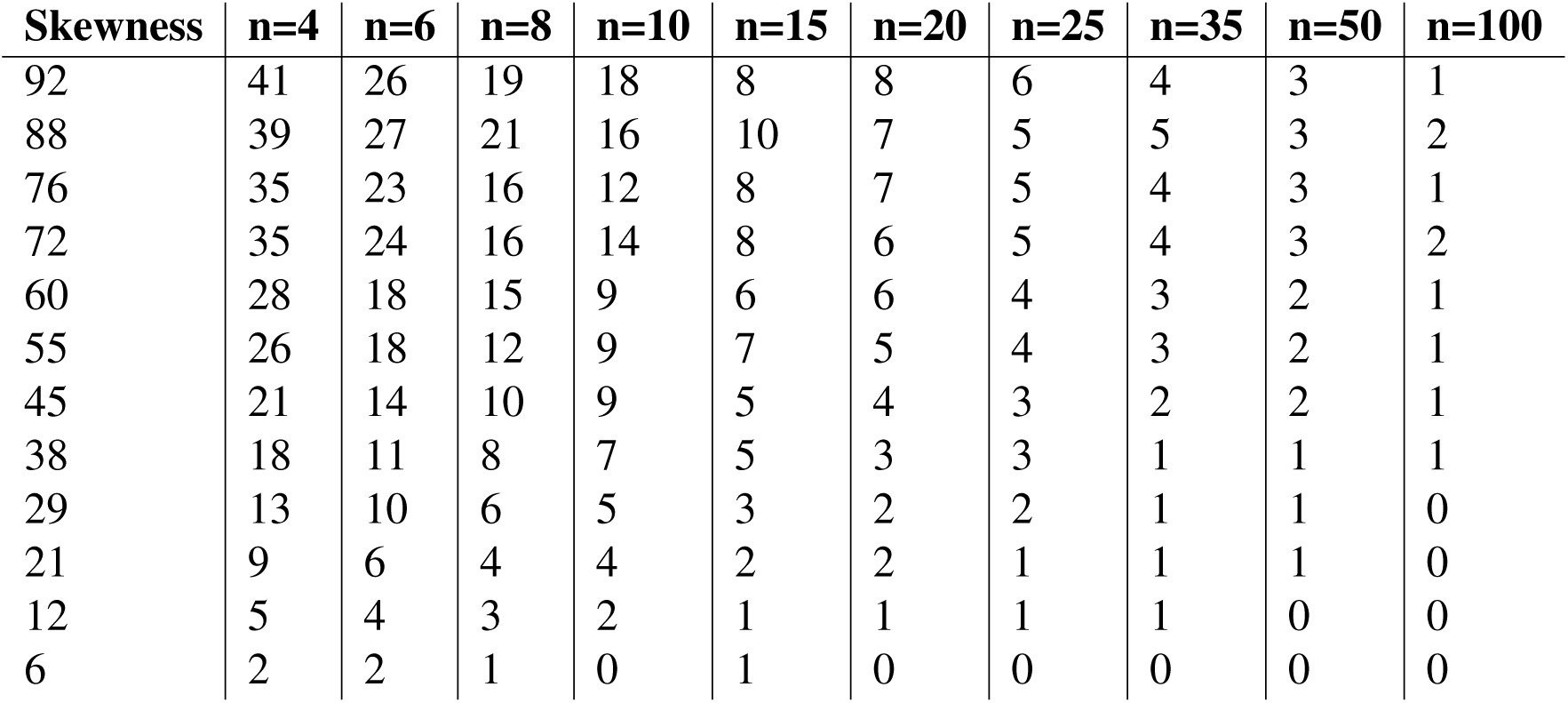
Bias estimation for Miller’s 12 ex-Gaussian distributions. Columns correspond to different sample sizes. Rows correspond to different distributions, sorted by skewness. There are several ways to define skewness using parametric and non-parametric methods. Following Miller 1988, and because the emphasis is on the contrast between the mean and the median in this article, we defined skewness as the difference between the mean and the median.

The results are also illustrated in Figure 3B. As reported by Miller (1988), bias can be quite large and it gets worse with decreasing sample sizes and increasing skewness. Based on these results, Miller (1988) made this recommendation:

> ’An important practical consequence of the bias in median reaction time is that sample medians must not be used to compare reaction times across experimental conditions when there are unequal numbers of trials in the conditions.’

According to Google Scholar, Miller (1988) has been cited 182 times as of the 8th of January 2019. A look at some of the oldest and most recent citations reveals that his advice has been followed. A popular review article on reaction times, cited 411 times, reiterates Miller’s recommendations (Whelan, 2008).

However, there are several problems with Miller’s advice, which we explore in the next sections, starting with one key omission from Miller’s assessment: the bias of the sample median can be corrected using a percentile bootstrap bias correction.

## Bias correction

A simple technique to estimate and correct sampling bias is the percentile bootstrap (Efron & Tibshirani, 1994). If we have a sample of n observations, here is how it works:

- sample with replacement n observations from the original sample
- compute the estimate (say the mean or the median)
- perform steps 1 and 2 *nboot* times
- compute the mean of the *nboot* bootstrap estimates

The difference between the estimate computed using the original sample and the mean of the *nboot* bootstrap estimates is a bootstrap estimate of bias.

To illustrate, let’s consider one random sample of 10 observations from the skewed distribution in Figure 1A, which has a population median of 508.7 (rounded to 509 in the figure):

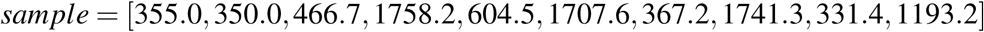

The median of the sample is 535.6, which over-estimates the population value of 508.7. Next, we sample with replacement 1,000 times from our sample, and compute 1,000 bootstrap estimates of the median. The distribution obtained is a bootstrap estimate of the sampling distribution of the median, as illustrated in Figure 4. The idea is this: if the bootstrap distribution approximates, on average, the shape of the sampling distribution of the median, then we can use the bootstrap distribution to estimate the bias and correct our sample estimate. However, as we’re going to see, this works on average, in the long-run. There is no guarantee for a single experiment.

**Figure 4.**
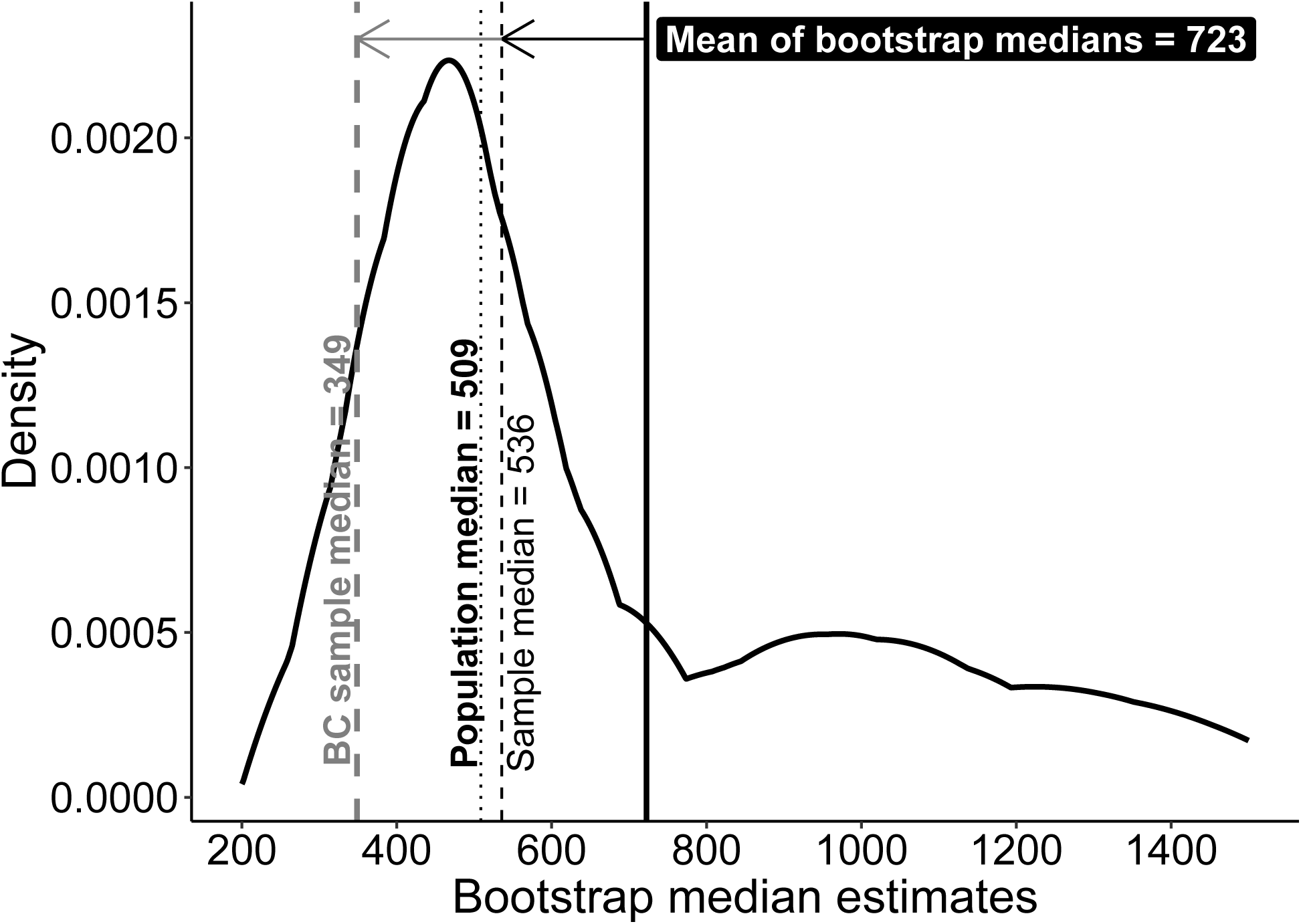
Bootstrap bias correction example: one experiment. The sample median (black dashed vertical line) overestimates the population value (black dotted vertical line). The kernel density estimate of 1,000 bootstrap estimates of the sample median suggests, correctly, that the median sampling distribution is positively skewed. The difference between the sample median and the mean of the bootstrap medians (thick black vertical line) defines the bootstrap estimate of the bias (black horizontal arrow). This estimate can be subtracted from the sample median (grey horizontal arrow) to obtain a bias corrected (BC) sample median (grey dashed vertical line).

The mean of the bootstrap estimates is 722.6. Therefore, our estimate of bias is the difference between the mean of the bootstrap estimates and the sample median, which is 187, as shown by the black horizontal arrow in Figure 4. To correct for bias, we subtract the bootstrap bias estimate from the sample estimate (grey horizontal arrow in Figure 4):

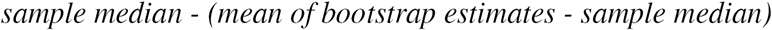

which is the same as:

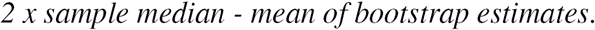

Here the bias corrected sample median is 348.6. So the sample bias has been reduced dramatically, clearly too much from the original 535.6. But bias is a long-run property of an estimator, so let’s look at a few more examples. We take 100 samples of n = 10, and compute a bias correction for each of them. The results of these 100 simulated experiments are shown in Figure 5. The arrows go from the sample median to the bias corrected sample median. The black vertical line shows the population median we’re trying to estimate.

**Figure 5.**
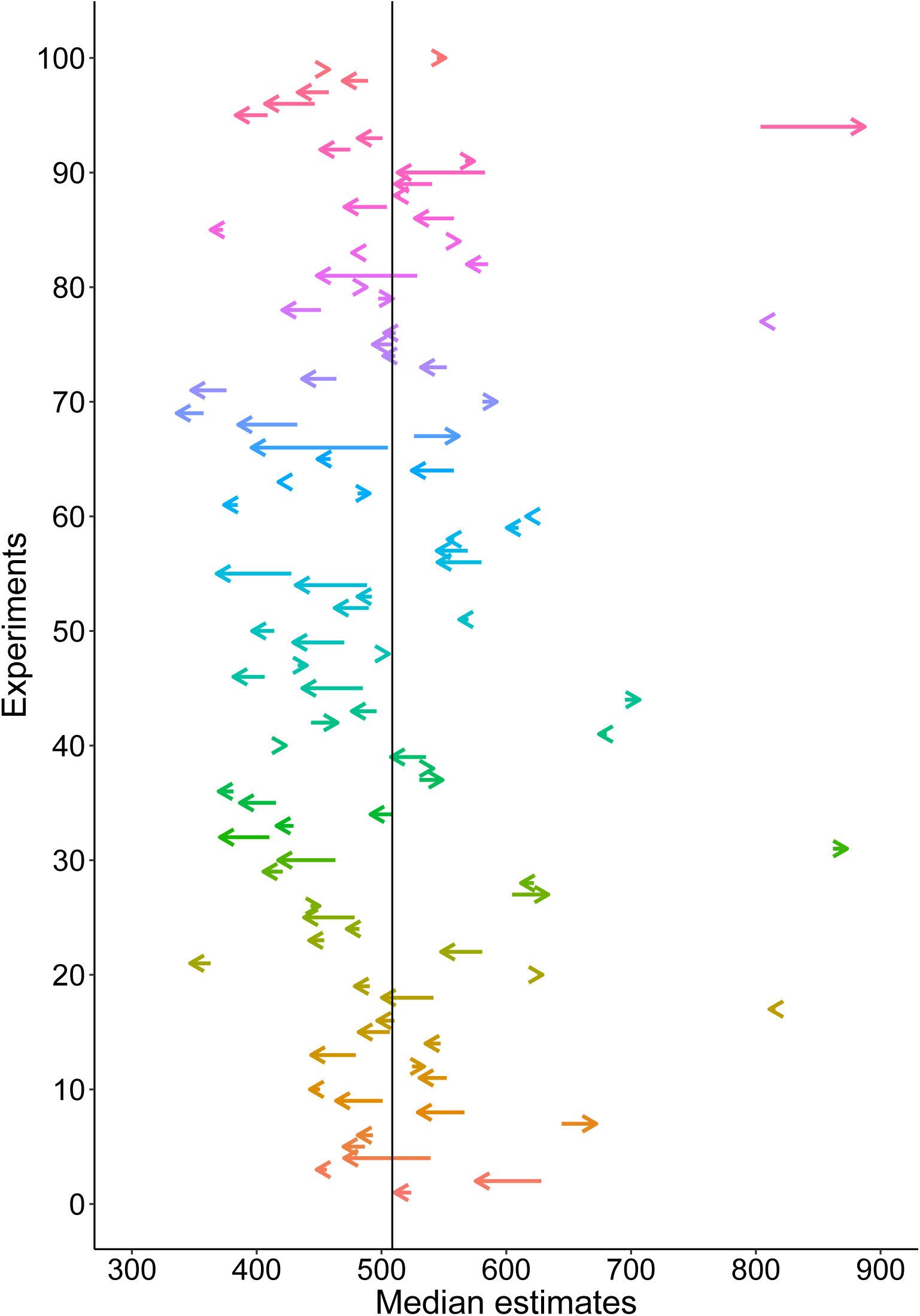
Bootstrap bias correction example: 100 experiments. For each experiment, 10 observations were sampled. Each arrow starts at the sample median for one experiment and ends at the bias corrected sample median. The bias was estimated using 200 bootstrap samples. The black vertical line marks the population median.

With *n* = 10, the sample estimates have large spread around the population and more so on the right than the left of the distribution. The bias correction also varies a lot in magnitude and direction, sometimes improving the estimates, sometimes making matters worse. Across experiments, it seems that the bias correction was too strong: the population median was 508.7, the average sample median was 515.1, but the average bias corrected sample median was only 498.8.

What happens if instead of 100 experiments, we perform 1000 experiments, each with *n* = 10, and compute a bias correction for each one? Now the average of the bias corrected median estimates is much closer to the true median: the population median was 508.7, the average sample median was 522.1, and the average bias corrected sample median was 508.6. So the bias correction works very well in the long-run for the median. But that’s not always the case: it depends on the estimator and on the amount of skewness (for instance, as we will see in the next section, bias correction fails for quantiles estimated using the Harrell-Davis estimator).

If we apply the bias correction technique to our median estimates of samples from Miller’s 12 distributions, we get the results in Figure 3C. For each iteration in the simulation, bias correction was performed using 200 bootstrap samples. The bias correction works very well on average, except for the smallest sample sizes. The failure of the bias correction for very small n is not surprising, because the shape of the sampling distribution cannot be properly estimated by the bootstrap from so few observations. More generally, we should keep in mind that the performance of bootstrap techniques depends on sample sizes and the number of resamples, among other factors (Efron & Tibshirani, 1994; Wilcox, 2017). From *n* = 10, the bias values are very close to those observed for the mean. So it seems that in the long-run, we can eliminate the bias of the sample median by using a simple bootstrap procedure. As we will see in another section, the bootstrap bias correction is also effective when comparing two groups. Before that, we need to look more closely at the sampling distributions of the mean and the median.

## Sampling distributions

The bias results presented so far rely on the standard definition of bias as the distance between the mean of the sampling distribution (here estimated using Monte-Carlo simulations) and the population value. However, using the mean to quantify bias assumes that the sampling distributions are symmetric and that we are only interested in the long-term performance of an estimator. If the sampling distributions are asymmetric and we want to know what bias to expect in a typical experiment, other measures of central tendency, such as the median, would characterise bias better than the mean.

Consider the sampling distributions of the mean and the median for different sample sizes from the ex-Gaussian distributions described in Figure 2. When skewness is low (6, first row of Figure 6), the sampling distributions are symmetric and centred on the population values: there is no bias. As we saw previously, with increasing sample size, variability decreases, which is why studies with larger samples provide more accurate estimations. The flip side is that studies with small samples are much noisier, which is why results across small n experiments can differ substantially (Button et al., 2013). By showing the variability across simulations, the sampling distributions also highlight an important aspect of the results: bias is a long-run property of an estimator; there is no guarantee that one value from a single experiment will be close to the population value, particularly for small sample sizes.

**Figure 6.**
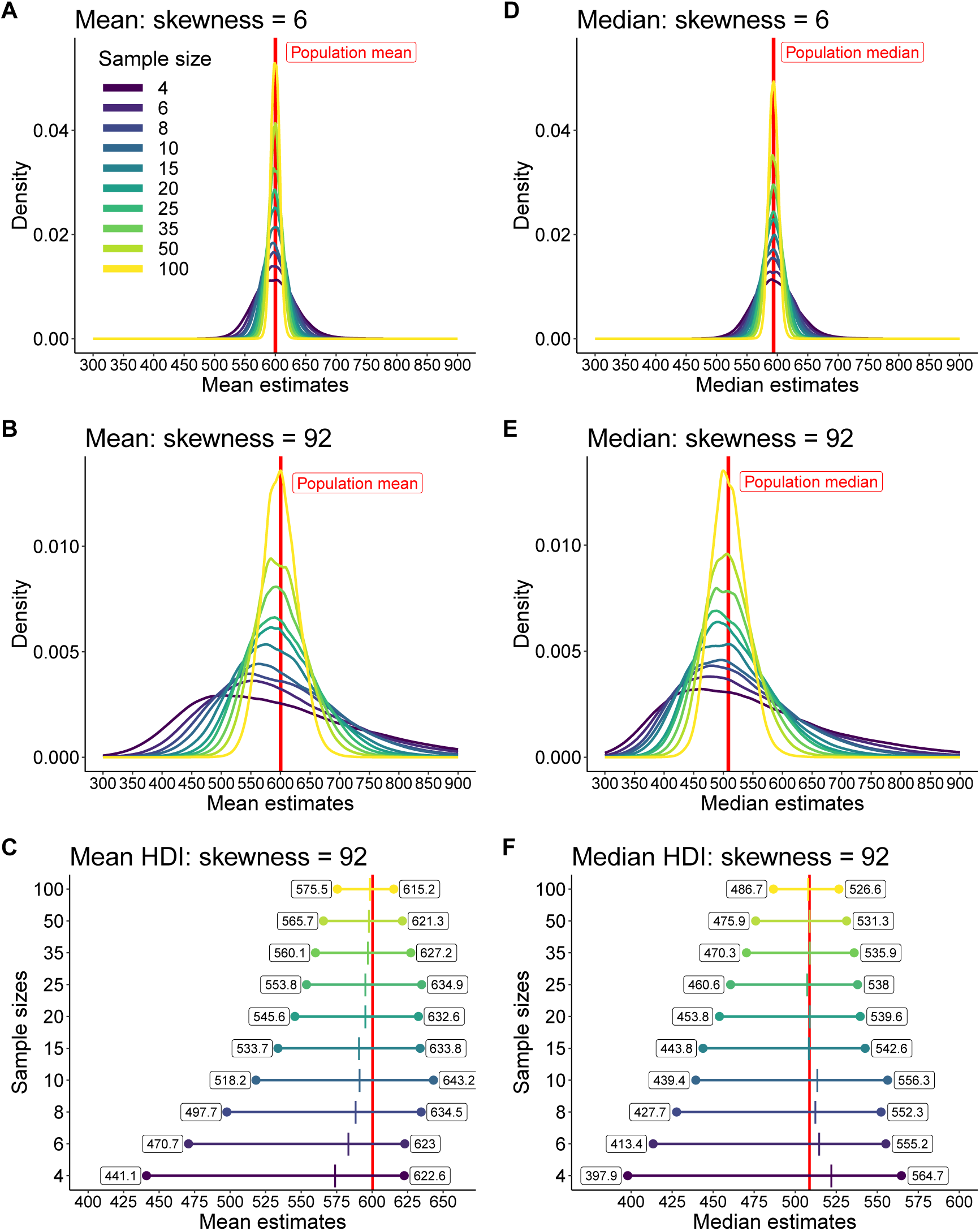
Sampling distributions of the mean and the median. In each panel, results for sample sizes from 4 to 100 are colour coded. The results are based on 10,000 samples for each sample size and skewness. Left column: results for the mean; right column: results for the median. The bottom row illustrates, for the mean (**C**) and the median (**F**), the 50% HDI of the distributions shown in the second row.

When skewness is large (92, second row of Figure 6), sampling distributions get more positively skewed with decreasing sample sizes. To better understand how the sampling distributions change with sample size, we turn to the last row of Figure 6, which shows 50% highest-density intervals (HDI). A HDI is the shortest interval that contains a certain percentage of observations from a distribution (Kruschke, 2013). For symmetric distributions, HDI and confidence intervals are similar, but for skewed distributions, HDI better capture the location of the bulk of the observations.Each horizontal line is a HDI for a particular sample size. The labels contain the values of the interval boundaries. The coloured vertical tick inside the interval marks the median of the distribution. The red vertical line spanning the entire plot is the population value.

For means and small sample sizes, the 50% HDI is offset to the left of the population mean, and so is the median of the sampling distribution. This demonstrates that the typical sample mean tends to under-estimate the population mean – that is to say, the mean sampling distribution is median biased. This offset reduces with increasing sample size, but is still present even for *n* = 100.

For medians and small sample sizes, there is a discrepancy between the 50% HDI, which is shifted to the left of the population median, and the median of the sampling distribution, which is shifted to the right of the population median. This contrasts with the results for the mean, and can be explained by differences in the shapes of the sampling distributions, in particular the larger skewness and kurtosis of the median sampling distribution compared to that of the mean. The offset between the sample median and the population value reduces quickly with increasing sample size. For *n* = 10, the median bias is already very small. From *n* = 15, the median sample distribution is not median bias, which means that the typical sample median is not biased.

Another representation of the sampling distributions is provided in Figure 7: 50% HDI of the biases are shown as a function of sample size. For both the mean and the median, the spread of the bias increases with increasing skewness and decreasing sample size. Skewness also increases the asymmetry of the bias distributions, but more so for the mean than the median.

**Figure 7.**
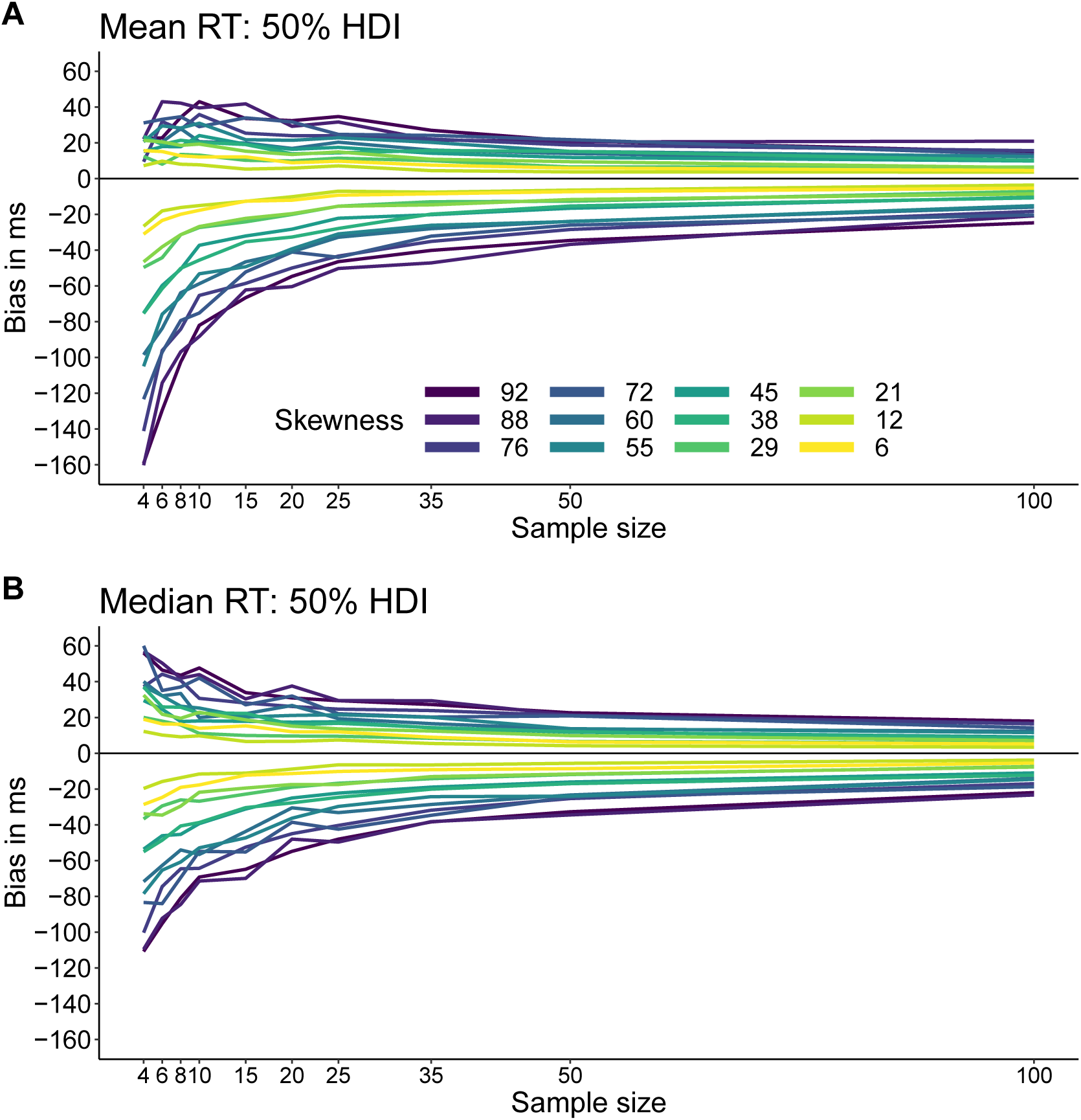
50% highest density intervals of the biases of the sample mean and the sample median as a function of sample size and skewness.

So is the mean also biased? According to the standard definition of bias, which is based on the distance between the population mean and the average of the sampling distribution of the mean, the mean is not biased. But this definition applies to the long-run, after we replicate the same experiment many times - 10,000 times in our simulations. So what happens in practice, when we perform only one experiment instead of 10,000? In that case, the median of the sampling distribution provides a better description of the typical experiment than the mean of the distribution. And the median of the sampling distribution of the mean is inferior to the population mean when sample size is small. So if we conduct one small n experiment and compute the mean of a skewed distribution, we’re likely to under-estimate the true value.

Is the median biased after all? The median is indeed biased according to the standard definition. However, with small n, the typical median (represented by the median of the sampling distribution of the median) is close to the population median, and the difference disappears for even relatively small sample sizes. In other words, in a typical experiment, the median shows very limited bias.

## Group differences: bias

Now that we better understand the sampling distributions of the mean and the median, we consider how these two estimators perform when we compare two groups. According to Miller (1988), because the median is biased when taking small samples from skewed distributions, group comparison can be affected if the two groups differ in sample size, such that real differences can be lowered or increased, and non-existent differences suggested. As a result, for unequal n, Miller advised against the use of the median. In this section, we put this advice to the test, by simulating experiments in which individual RT distributions are summarised using the mean or the median, and these individual means and medians are then compared across participants - a typical hierarchical experimental design. We consider cases in which an effect does not exist, to look at false positives, and cases in which an effect does exist, to look at true positives.

We assessed the problem using a simulation in which we drew samples from populations defined by the same 12 distributions used by Miller (1988), as described previously. Group 2 had a constant size of *n* = 200; group 1 had size *n* = 10 to *n* = 200, in increments of 10. For the mean, the results of 10,000 iterations are presented in Figure 8A. All the bias values are near zero, as expected.

**Figure 8.**
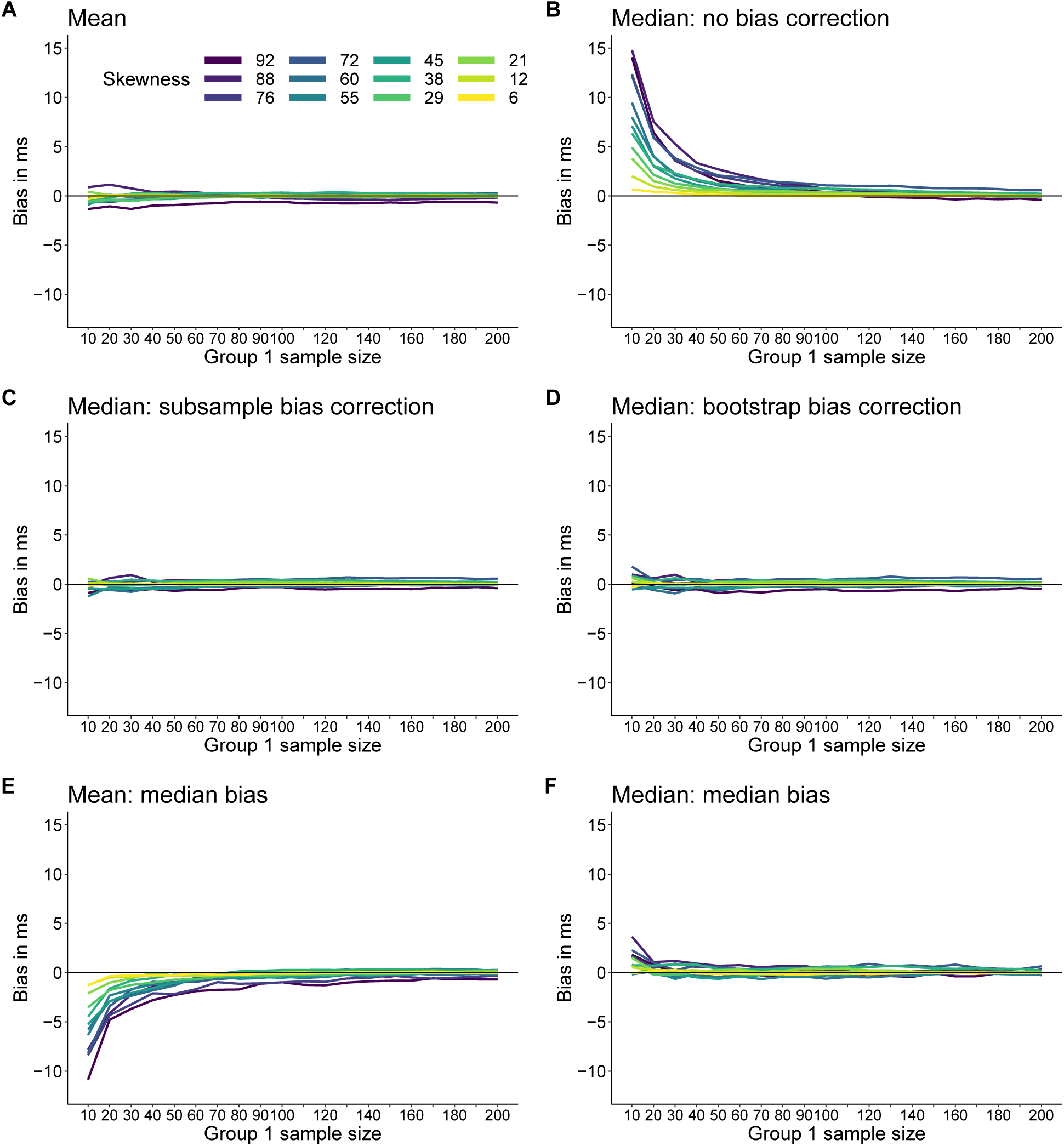
Bias estimation for the difference between two independent groups. Group 2 is always sampled from the least skewed distribution and has size n = 200. The size of group 1 is indicated along the x axis in each panel. Group 1 is sampled from Miller’s 12 skewed distributions and the results are colour coded by skewness. **A.** Bias for mean reaction times. **B.** Bias for median reaction times. **C.** Bias for median reaction times after subsample bias correction. **D.** Bias for median reaction times after bootstrap bias correction. **E.** Median bias for mean reaction times. **F.** Median bias for median reaction times.

**Figure 9.**
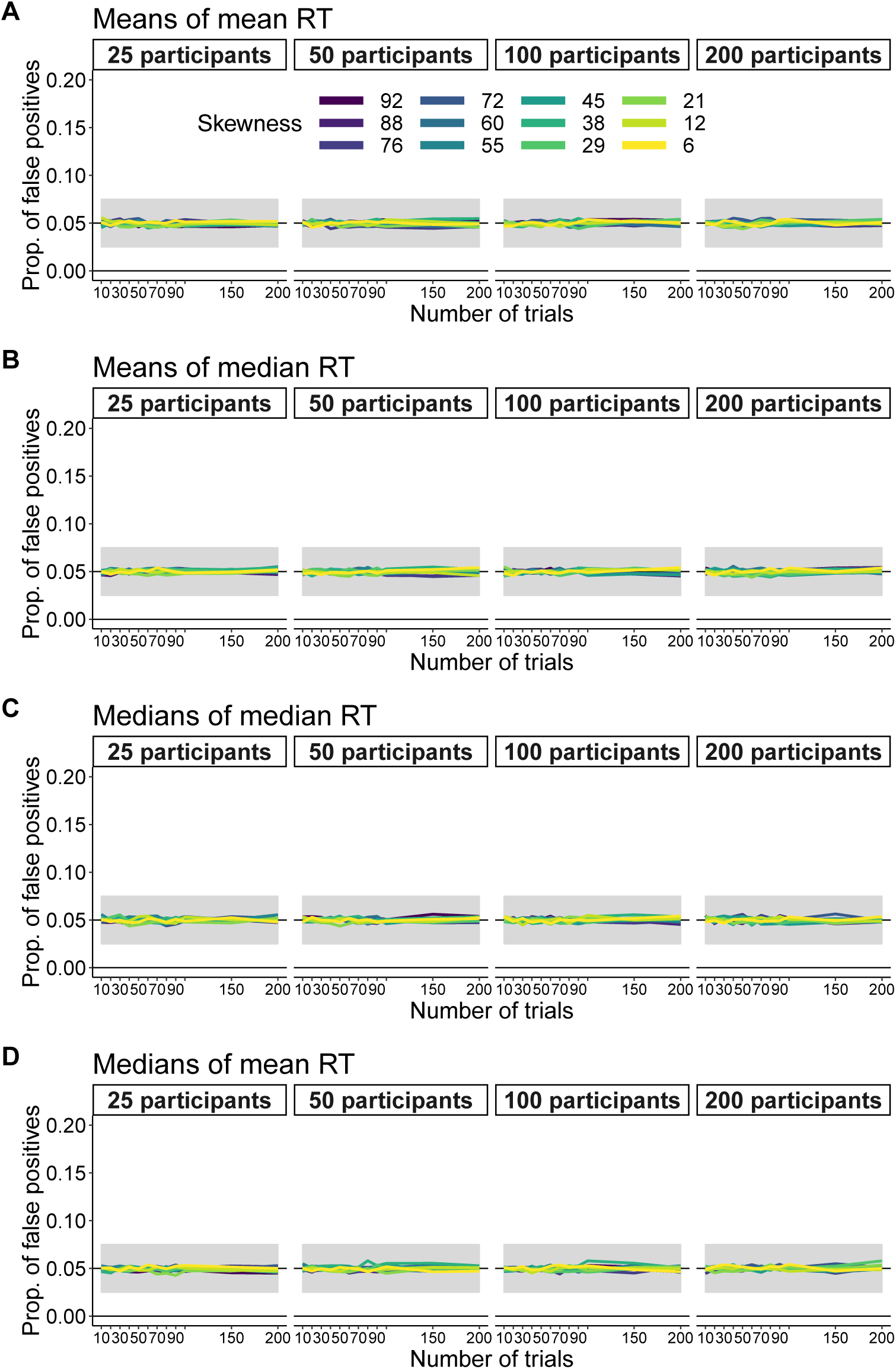
False positives: equal number of trials. Results are shown for group means of differences between individual mean reaction times **A**, means of medians **B**, medians of medians **C** and medians of means **D**. The shaded grey areas represent the minimally satisfactory range suggested by Bradley (1978): 0.025-0.075. The black horizontal lines behind the shaded areas mark the expected 0.05 proportion of false positives.

Results for the median are presented in Figure 8B. Bias increases with skewness and sample size difference (the difference gets larger as the sample size of group 1 gets smaller). At least about 90-100 trials in Group 1 are required to bring bias to values similar to the mean.

Next, let’s find out if we can correct the bias. Bias correction was performed in 2 ways: with the bootstrap, as explained in the previous section, and with a different approach using subsamples. The second approach was suggested by Miller (1988):

> ’Although it is computationally quite tedious, there is a way to use medians to reduce the effects of outliers without introducing a bias dependent on sample size. One uses the regular median from Condition F and compares it with a special “average median” (Am) from Condition M. To compute Am, one would take from Condition M all the possible subsamples of Size f where f is the number of trials in Condition F. For each subsample one computes the subsample median. Then, Am is the average, across all possible subsamples, of the subsample medians. This procedure does not introduce bias, because all medians are computed on the basis of the same sample (subsample) size.’

Using all possible subsamples would take far too long; for instance, if one group has 5 observations and the other group has 20 observations, there are 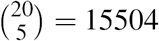 subsamples to consider. Slightly larger sample sizes would force us to consider millions of subsamples. So instead we computed K random subsamples. We arbitrarily set K to 1,000. Although this is not what Miller (1988) suggested, this shortcut should reduce bias to some extent if it is due to sample size differences. The results are presented in Figure 8C and show that the K loop approach works very well. But we need to keep in mind that it works in the long-run: for a single experiment there is no guarantee, similarly to the bootstrap procedure. The bias of the group differences between medians can also be handled by the bootstrap. Bias correction using 200 bootstrap samples for each simulation iteration leads to the results in Figure 8D: overall the bootstrap bias correction works very well. At most, for *n* = 10, the median’s maximum bias across distributions is 1.79 ms, whereas the mean’s is 0.88 ms.

So in the long-run, the bias in the estimation of differences between medians can be eliminated using the subsampling or the percentile bootstrap approaches. Because of the skewness of the sampling distributions, we also consider the median bias: the bias observed in a typical experiment. In that case, the difference between group means tends to underestimate the population difference, as shown in Figure 8E. For the median, the median bias is much lower than the standard (mean) bias, and near zero from *n* = 20 (Figure 8F). Thus, for a typical experiment, the difference between group medians actually suffers less from bias than the difference between group means. Also note that if the median is adjusted to be unbiased, a consequence is that it is now median biased.

Based on our results so far, we conclude that Miller’s (1988) advice was inappropriate because, when comparing two groups, bias in a typical experiment is actually negligible. To be cautious, when sample size is relatively small, it could be useful to report median effects with and without bootstrap bias correction. It would be even better to run simulations to determine the sample sizes required to achieve an acceptable measurement precision, irrespective of the estimator used (Schönbrodt & Perugini, 2013; Peters & Crutzen, 2017; Rothman & Greenland, 2018). We provide an example of such simulation of measurement precision in a later section. Next, building up from what we have learned so far, we turn to a more complicated situation, in which skewness is considered at two levels of analysis: for each participant/condition, and at the group level.

## Group differences: error rates

When researchers compare groups, bias and sampling distributions are, regrettably, not at the top of their minds. Most researchers are interested in false positives (type I errors) and true positives (statistical power) associated with a test. So in this section, we extend the previous simulations to consider the error rates associated with tests using the mean and the median in a hierarchical setting involving a within-subject (repeated-measure) design: single-trials are sampled from 2 conditions in multiple participants, each individual distribution of trials is summarised, and the individual summaries are compared across participants. For instance, when means are employed, a standard approach is to compare means of means: that is, for each condition and each participant, the distribution is summarised using the mean; then the means from the two conditions are subtracted; finally, the individual differences between means are assessed across participants by computing a one-sample t-test on group means. We simulated this situation using the 12 ex-Gaussian distributions used so far, 10,000 iterations, an arbitrary alpha level of 0.05, and different numbers of trials and participants. In the case of an equal number of trials in each condition, the results show false positives close to the nominal level (0.05) for all skewness levels (Figure 9A). Although computing group means of individual differences between means is a popular choice, the choice of estimator must be made at the two levels of analysis. If we restrict our options to the mean and the median, we can consider three more combinations, for which results similar to that of the mean were obtained: group means of individual differences between medians, medians of medians, medians of means (Figure 9B-D). Tests on medians were performed using the method by Hettmansperger and Sheather (1986).

What happens if we use unequal sample sizes between conditions, as done in the previous section? For means of means, the false positives are again at the nominal level (Figure 10A). However, the results change drastically for group means of individual medians: with small sample sizes, the proportion of false positives increases with skewness and with the number of participants (Figure 10B). With *n* = 10 or 20, 200 participants and very skewed distributions, false positives are committed more than 50% of the time! This unfortunate behaviour is due to the (mean) bias of the sample median.

**Figure 10.**
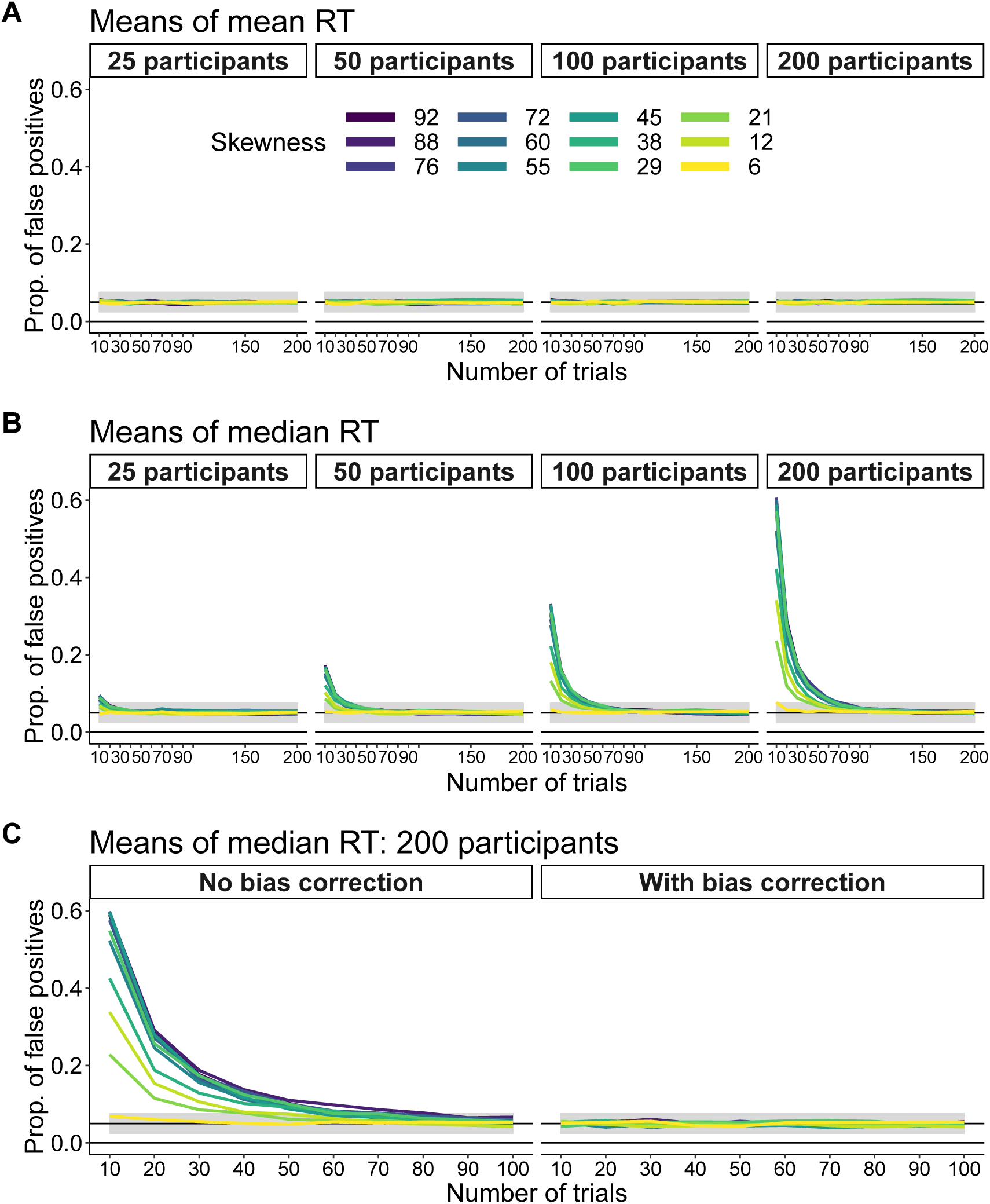
False positives: group means and unequal numbers of trials. **A.** Bias for group means of indvidual mean reaction times. **B.** Bias for group means of individual median reaction times. **C.** Effect of the percentile bootstrap bias correction on inferences based on the means of median reaction times calculated for 200 participants. The left panel is a replication of the right most panel from B, using 2,000 iterations instead of 10,000. In the right panel, a bias correction with 200 bootstrap samples was applied.

The effect of the bias of the sample median can be addressed by increasing the number of trials (Figure 10B). The proportion of false positives can also be brought close to the nominal level by using the percentile bootstrap bias correction (Figure 10C). Yet another strategy is to assess group medians of individual medians instead of group means (Figure 11A). Indeed, because the sample median is not median biased, computing medians of medians gets rid of the bias. Conversely, because the sample mean is median biased, performing tests on medians of means for small sample sizes leads to an increase in false positives with increasing skewness and number of participants (Figure 11B).

**Figure 11.**
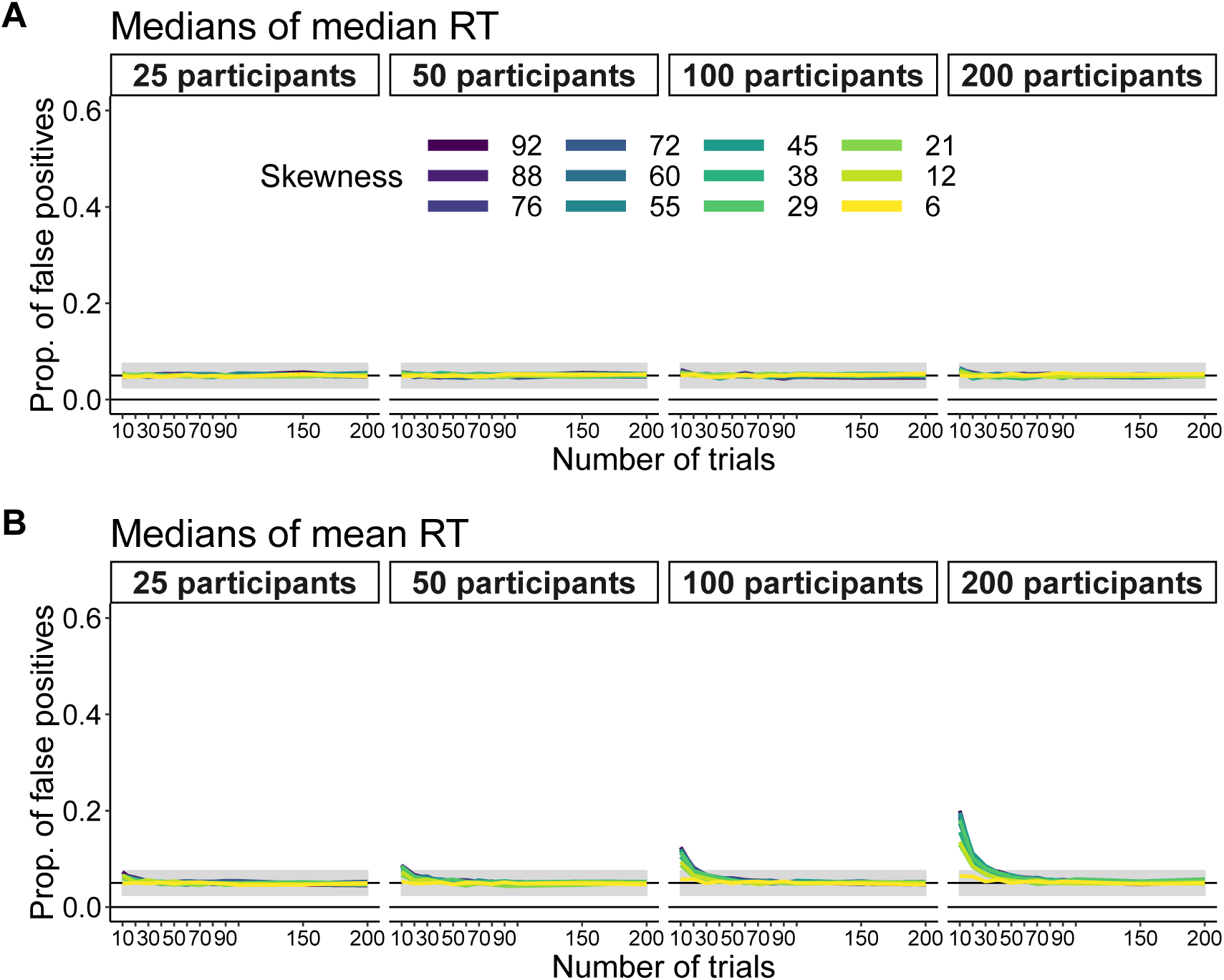
False positives: group medians and unequal numbers of trials. **A.** Bias for medians of median reaction times. **B.** Bias for medians of mean reaction times.

### Effects of asymmetry and outliers on error rates

From the previous results, we could conclude that, if the goal is to keep false positives at the nominal level, using group means of individual means or group medians of individual medians are satisfactory options in the long-run. However, the choice of estimators used to make group inferences depends on the shape of the group distributions. In particular, skewness and outliers can have very detrimental effects on the performance of certain test statistics, such as t-tests on means. We illustrate this problem with a simulation with 10,000 iterations using *g*&*h* distributions: the median of these distributions is always zero, *g* controls the asymmetry of the distribution and *h* controls the thickness of the tails (Hoaglin, 1985a). With increasing *h*, outliers are more and more frequent. Let’s first consider false positives as a function of the *g* parameter (Figure 12). For simplicity, we only simulate what happens at the group level: the values considered could be any type of differences between means, medians or any other quantities.

**Figure 12.**
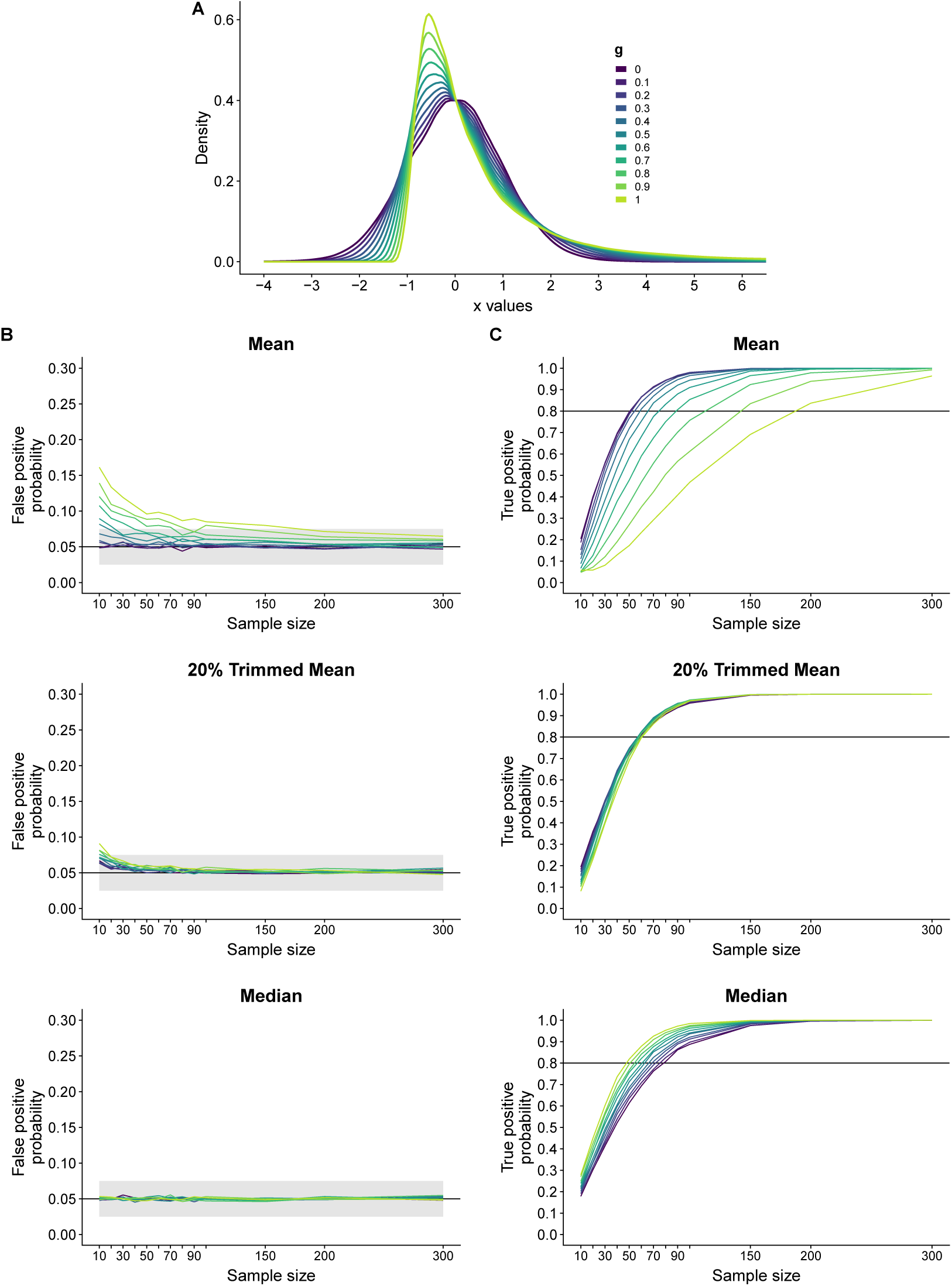
Group inferences using *g*&*h* distributions: false and true positives as a function of the *g* parameter. **A.** Illustrations of probability density functions for *g* varying from 0 to 1, *h* = 0. With *g* = 1 and *h* = 0, the distribution has the same shape as a lognormal distribution. **B.** False positive results for the mean, the 20% trimmed mean and the median. Samples were drawn from populations with means, 20% trimmed means and medians of 0. **C.** True positive results (power) for the mean, the 20% trimmed mean and the median. An effect was simulated by a constant to samples drawn from populations with means, 20% trimmed means and medians of 0.

As expected, for normal distributions (*g* = 0), the false positives are at the nominal level (5%). However, with increasing g, the probability of false positives increases, and much more so with smaller sample sizes. In contrast, tests on medians are unaffected by skewness. A third estimator, the 20% trimmed mean, gives intermediate results: it is affected by skewness much less than the mean and only for the smallest sample sizes. In a 20% trimmed mean, observations are sorted, 20% of observations are eliminated from each end of the distribution and the remaining observations are averaged. Inferences based on the 20% trimmed mean were made via an extension of the one-sample t-test proposed by Tukey and McLaughlin (1963). Means and medians are special cases of trimmed means: the mean corresponds to zero trimming and the median to 50% trimming.

What happens when we keep *g* constant (*g* = 0.3) and *h* varies (Figure 13A)? In the case of t-tests on means, for *h* = 0, false positive probabilities are near the nominal level, but then increase with *h* irrespective of sample size (Figure 13B). In contrast, medians are not affected by *h*, and for 20% trimmed means, false positives increase only when *n* = 10. When *g* = 0 and h varies, the proportion of false positives for the mean decreases slightly below 0.05 irrespective of sample size, whereas tests on 20% trimmed mean and median are unaffected (see supplementary notebook *sim gp g&h*).

**Figure 13.**
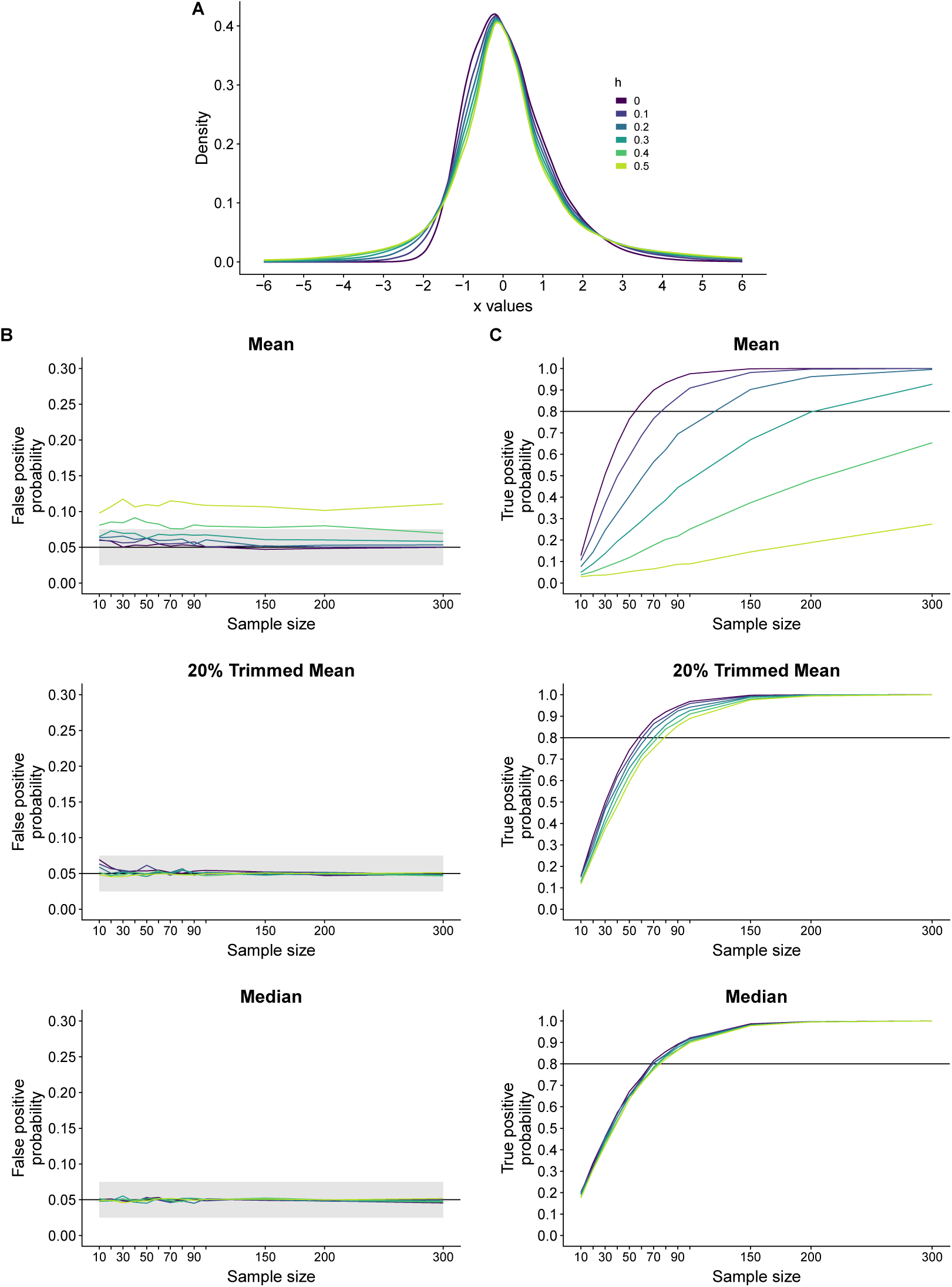
Group inferences using g & h distributions: false and true positives as a function of the h parameter. **A.** Illustrations of probability density functions for h varying from 0 to 0.5, *g* = 0.3. **B.** False positive results for the mean, the 20% trimmed mean and the median. **C.** True positive results (power) for the mean, the 20% trimmed mean and the median.

When it comes to true positives (power), t-tests on means perform well for low levels of asymmetry. However, with increasing asymmetry, power can be substantially affected (Figure 12C; an effect was created by shifting each distribution by a constant). Power with 20% trimmed means shows very little effects of asymmetry, except for the smallest sample sizes, but much less so than the mean. The median is associated with a seemingly odd behaviour: power increases with asymmetry! Also, power is higher for the mean when *g* = 0, a well-known result due to the over-estimation of the standard-error of the median under normality (Wilcox, 2017). Extra illustrations are available in notebook *samp dist*, showing that, for the ex-Gaussian distributions considered here, the sampling distribution of the mean becomes more variable than that of the median with increasing skewness and lower sample sizes.

When the probability of outliers increases (*h* increases), the power of t-tests on means can be strongly affected (Figure 13C), t-tests on 20% trimmed means much less so, and tests on medians almost not at all. Altogether, the *g*&*h* simulations demonstrate that the choice of estimator can have a strong influence on long-run error rates: t-tests on means are very sensitive to the asymmetry of distributions and the presence of outliers. But the results should not be used to advocate the use of medians in all situations - as we will see, no method dominates and the best approach depends on the empirical question and the type of effect. Certainly, relying blindly on the mean in all situations is unwise.

## Group differences: power curves in different situations

With the simulations in the previous section, we explored false positives and true positives at the group level, depending on the shape of the distributions of differences. For true positives, we considered a simple example in which the distribution of differences is shifted by a constant. However, distributions can differ in various ways. So here we assess power in different situations in a hierarchical setting: varying numbers of trials are sampled from an ex-Gaussian distribution in 2 conditions, in varying numbers of participants. These simulations assume no between-participant variability. We will provide a more realistic simulation later. First, we consider a uniform shift. In condition 1, the ex-Gaussian parameters were *mu* = 500, *sigma* = 50 and *tau* = 200. Parameters in condition 2 were the same, but each sample was shifted by 20. For illustration, Figure 14A shows two distributions with each *n* = 1000 and a shift of 50.

**Figure 14.**
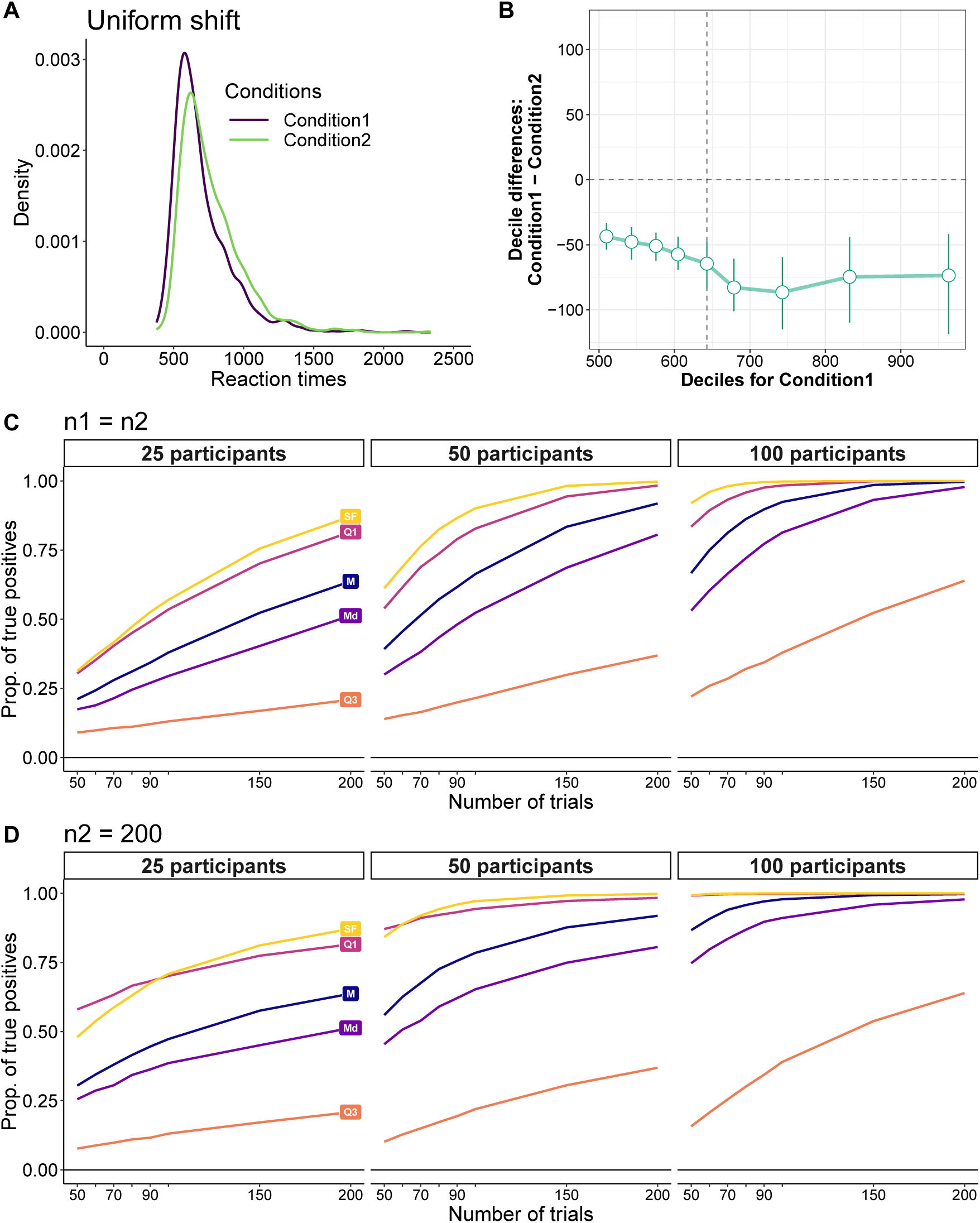
True positives: uniform shift. **A.** Example sampling distributions in conditions 1 and 2. **B.** Shift function. The differences between deciles are plotted as a function of deciles in condition 1. Deciles were computed using the Harrell-Davis quantile estimator. The error bars show 95% confidence intervals computed using a percentile bootstrap. **C.** Proportion of true positives (power) for equal sample sizes. **D.** Power results for unequal sample sizes. M = mean, Md = median, Q1 = first quartile, Q3 = third quartile, SF = hierarchical shift function.

To better understand how the distributions differ, panel B provides a shift function, in which the difference between the deciles of the two conditions are plotted as a function of the deciles in condition 1 - see details in Rousselet et al. (2017). The decile differences are all negative, showing stochastic dominance of condition 2 over condition 1. The function is not flat because of random sampling and limited sample size. Power curves are shown in panels C and D, respectively for equal and unequal sample sizes. Naturally, power increases with the number of trials and the number of participants. And for all combinations of parameters the mean is associated with higher power than the median.

In our example, other quantities are even more sensitive than the mean to distribution differences. For instance, considering that the median is the second quartile, looking at the other quartiles can be of theoretical interest to investigate effects in early or later parts of distributions. This could be done in several ways - here we performed tests on the group median of the individual quartile differences. For the uniform shift considered here, inferences on the third quartile (Q3) lead to lower power than the median. Using the first quartile (Q1) instead leads to a large increase in power relative to the mean. Q1 better captures uniform shifts because the early part of skewed distributions is much less variable than other parts, leading to more consistent differences across samples. Even with *n* = 1000, this lower variability can be seen in Figure 14B. So we might be able to achieve even higher power by considering even smaller quantiles than the first quartile, which could be motivated by an interest in the fastest responses (Rousselet, Macé & Fabre-Thorpe, 2003; Bieniek, Bennett, Sekuler & Rousselet, 2016; Reingold & Sheridan, 2018). But if the goal is to detect differences anywhere in the distributions, a more systematic approach consists in quantifying differences at multiple quantiles (K. Doksum, 1974; K. A. Doksum & Sievers, 1976).

### The hierarchical shift function

There is a rich tradition of using quantile estimation to understand the shape of distributions and how they differ (K. A. Doksum & Sievers, 1976; Hoaglin, 1985b; Marden et al., 2004), in particular to compare RT distributions (De Jong, Liang & Lauber, 1994; Pratte et al., 2010; Balota & Yap, 2011; Rousselet et al., 2017; Ellinghaus & Miller, 2018). Here we consider the case of the deciles, but other quantiles could be used. First, for each participant and each condition, the sample deciles are computed over trials. Second, for each participant, condition 2 deciles are subtracted from condition 1 deciles - we’re dealing with a within-subject (repeated-measure) design. Third, for each decile, the distribution of differences is subjected to a one-sample test. Fourth, a correction for multiple comparisons is applied across the 9 one-sample tests. We call this procedure a hierarchical shift function. There are many options available to implement this procedure and the example used here is not the definitive answer: the goal is simply to demonstrate that a relatively simple procedure can be much more powerful and informative than standard approaches.

In creating a hierarchical shift function we need to make three choices: a quantile estimator, a statistical test to assess quantile differences across participants, and a correction for multiple comparisons technique. The deciles were estimated using algorithm 8 described in (Hyndman & Fan, 1996). This estimator has the advantage, like the median, of being median unbiased and its standard (mean) bias can be corrected using the bias bootstrap correction (see detailed simulations in supplementary notebooks *hd bias* and *sf bias*). The main limitation of this estimator (and other estimators relying on the weighted average of one or two order statistics) is its poor handling of tied values. Tied values are not expected for continuous distributions such as RT, unless the raw data are rounded. If tied values are likely to occur, a good option is the Harrell-Davis quantile estimator (Harrell & Davis, 1982), which performs well in conjunction with the percentile bootstrap (Wilcox & Erceg-Hurn, 2012; Wilcox, Erceg-Hurn, Clark & Carlson, 2014). However, in the situations considered here, the Harrell-Davis estimator is both mean and median biased, and the bias bootstrap correction has limited effects on this bias (notebooks *hd bias* and *sf bias*). Again, no methods dominate.

The group comparisons were performed using a one-sample t-test on 20% trimmed means. In simulations, the 20% trimmed mean gave false positive rates closest to the nominal value relative to the mean, the median or the 10% trimmed mean (see simulation results in notebook *sim gp fp*). The four methods had similar power, with the mean performing the best, followed by the trimmed means and last the median (see simulation results in notebook *sim gp tp*). Given that our simulations did not include outliers, to which the mean is very sensitive, it seems safer to use the 20% trimmed mean by default - a choice consistent with extant results (Wilcox, 2017). The correction for multiple comparisons employed Hochberg’s strategy (Hochberg, 1988), which guarantees that the probability of at least one false positive will not exceed the nominal level as long as the nominal level is not exceeded for each quantile (Wilcox et al., 2014); the same strategy is applied in other shift function applications (Wilcox, 2017). The power curves for the hierarchical shift function are labelled SF in Figure 14C and D. SF’s power curves are very similar to those obtained for Q1. The advantage of SF is that the location of the distribution difference can be interrogated, which is impossible if inferences are limited to a single quantile such as Q1.

SF and Q1 methods are also very similar in their sensitivity to early differences illustrated in Figure 15, and much more so than methods based on the mean, the median, or Q3. These early differences were created by sampling from two ex-Gaussian distributions with parameters *mu* = 500, *sigma* = 50 and *tau* = 200. Condition 1 was centred on its third quartile, multiplied by 1.1 to stretch it, and then shifted by its third quartile. Early differences are well documented in the Simon task, for which the two conditions differ mostly in the fastest reaction times, and these differences progressively weaken and sometimes reverse sign (De Jong et al., 1994; Pratte et al., 2010; Schwarz & Miller, 2012). In our example, the median dominates the mean in terms of power in all situations.

**Figure 15.**
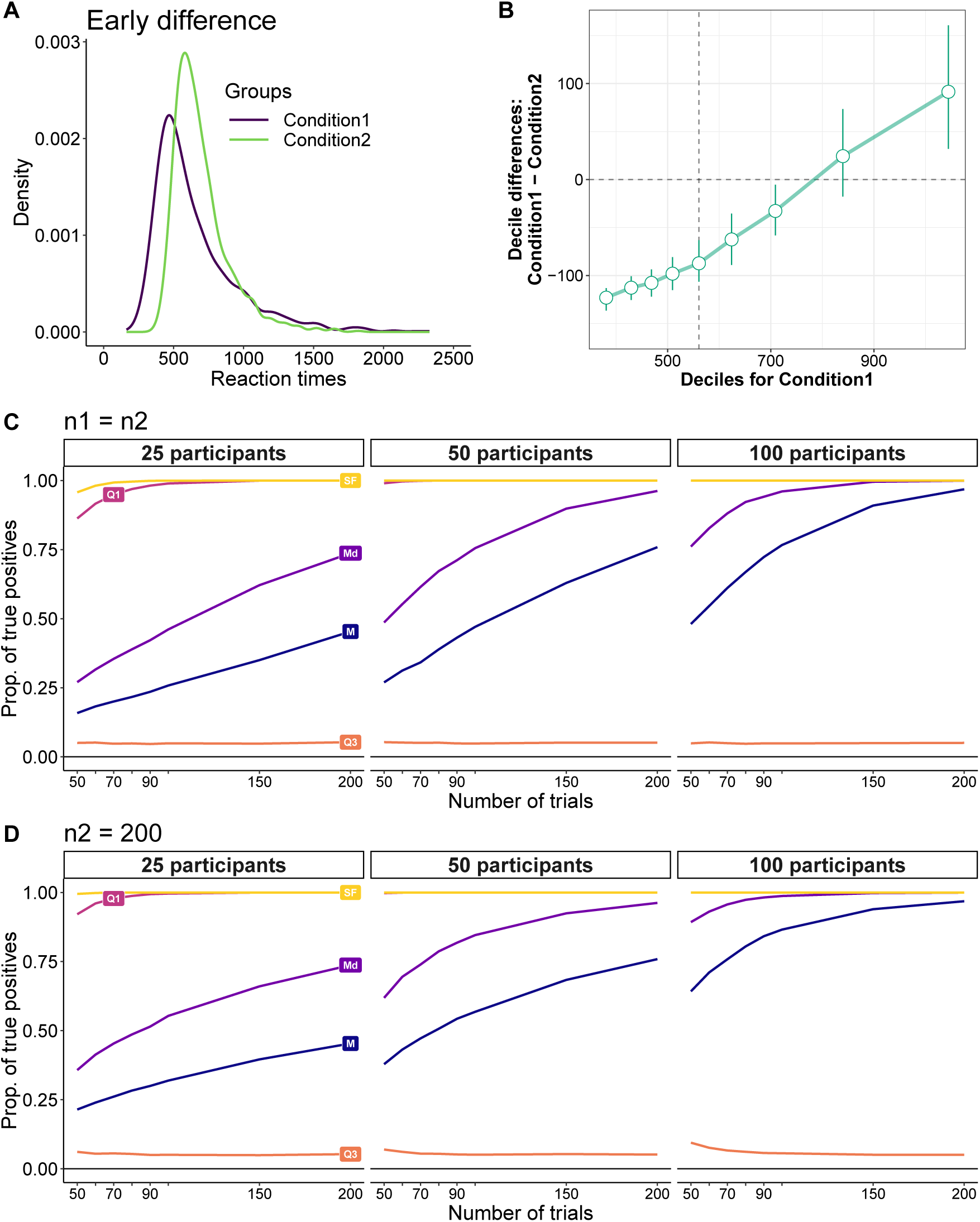
True positives: early difference. **A.** Example sampling distributions in conditions 1 and 2. For illustration, this distribution was created as explained in the text, but with a multiplication by 1.5 to create a stronger stretch. **B.** Shift function. **C.** Proportion of true positives (power) for equal sample sizes.**D.** Power results for unequal sample sizes.

Finally, we consider a situation in which a late effect is observed Figure 16. This situation was modelled by two ex-Gaussian distributions with the same parameters *mu* = 500 and *sigma* = 50 but different tau parameters, one set to 200, another one set to 215. In that situation, the mean dominates all methods, including SF as well as performing one-sample t-tests on the mean of differences in tau estimates obtained by fitting ex-Gaussian distributions to the data using the maximum likelihood method (Massidda, 2013). Thus, in this particular situation, the median performs poorly and the strong sensitivity of the mean to the right tail of the distribution is particularly useful, at least providing outliers do not affect group assessment.

**Figure 16.**
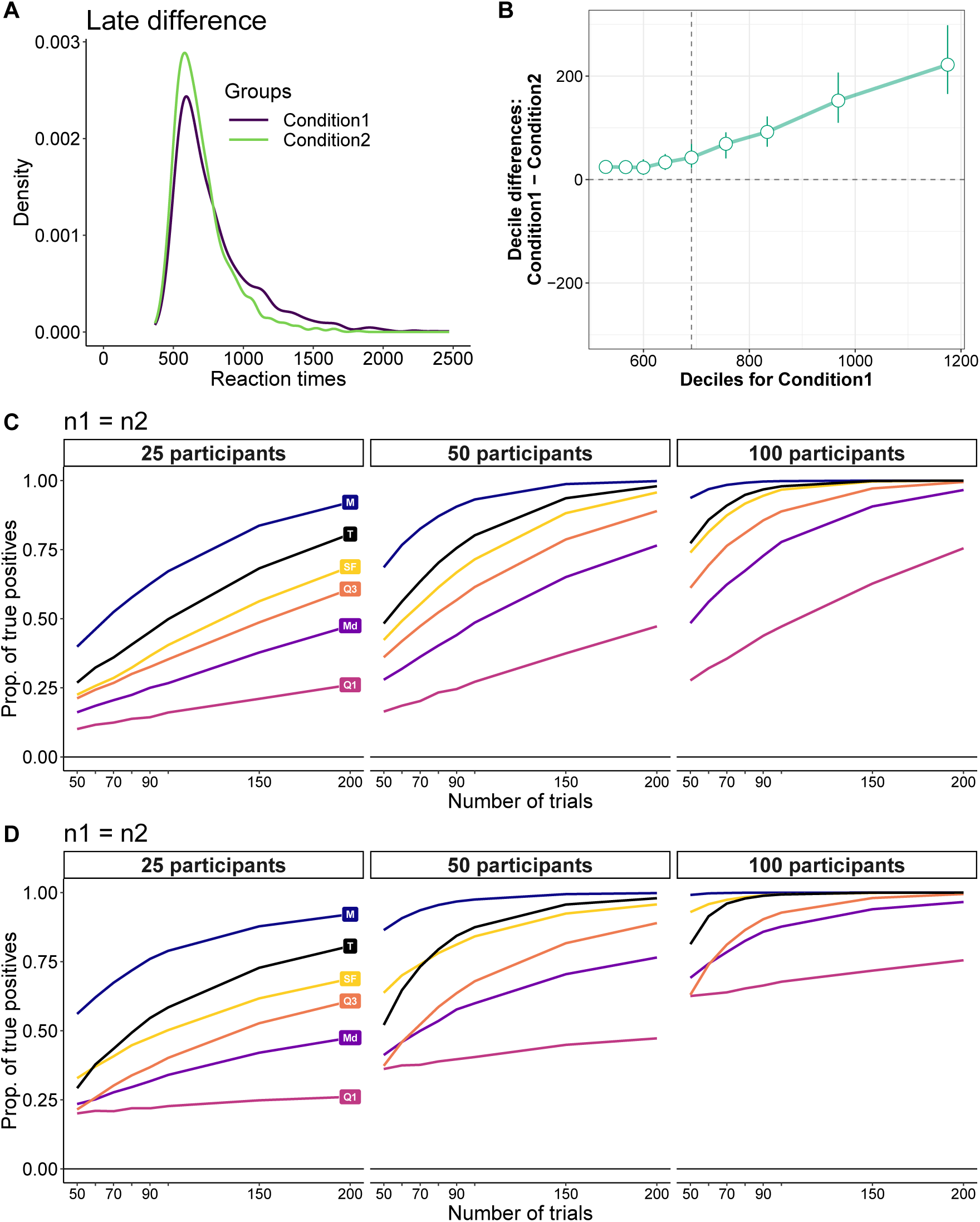
True positives: late difference. **A.** For illustration, the difference in the tau component was exaggerated. The two distributions had mu=500 and sigma=50, but tau was 200 in condition 2 and 300 in condition 1. **B.** Shift function. **C.** Proportion of true positives (power) for equal sample sizes. **D.** Power results for unequal sample sizes. T = one-sample t-tests on differences between the means of the tau component estimated using an ex-Gaussian fit to the data.

In summary, the examples considered so far demonstrate that that no method dominates: different methods better handle different situations. Crucially, neither the mean nor the median are sufficient or even necessary to compare skewed distributions. Better tools are available; in particular, considering multiple quantiles of the distributions allow us to get a deeper understanding of how distributions differ. This can easily be done using the R functions in the reproducibility package for this article (Rousselet & Wilcox, 2018a) and in a recent review (Rousselet et al., 2017).

## Applications to a large dataset

The simulations above suggest that using the median is appropriate in many situations involving skewed distributions and is preferable to the mean in some situations. In this final section we consider what happens when we deal with real RT distributions instead of simulated ones, as we have done so far. To find out, we look at the behaviour of the mean, the median and the hierarchical shift function in a large dataset of reaction times from participants engaged in a lexical decision task. The data are from the French lexicon project (FLP) (Ferrand et al., 2010). After removing a few participants who did not pay attention to the task (very low accuracy or too many late responses), we’re left with 959 participants. Each participant had between 996 and 1001 trials for each of two conditions, Word and Non-Word. Figure 17 illustrates reaction time distributions from 100 randomly sampled participants in the Word and Non-Word conditions.

**Figure 17.**
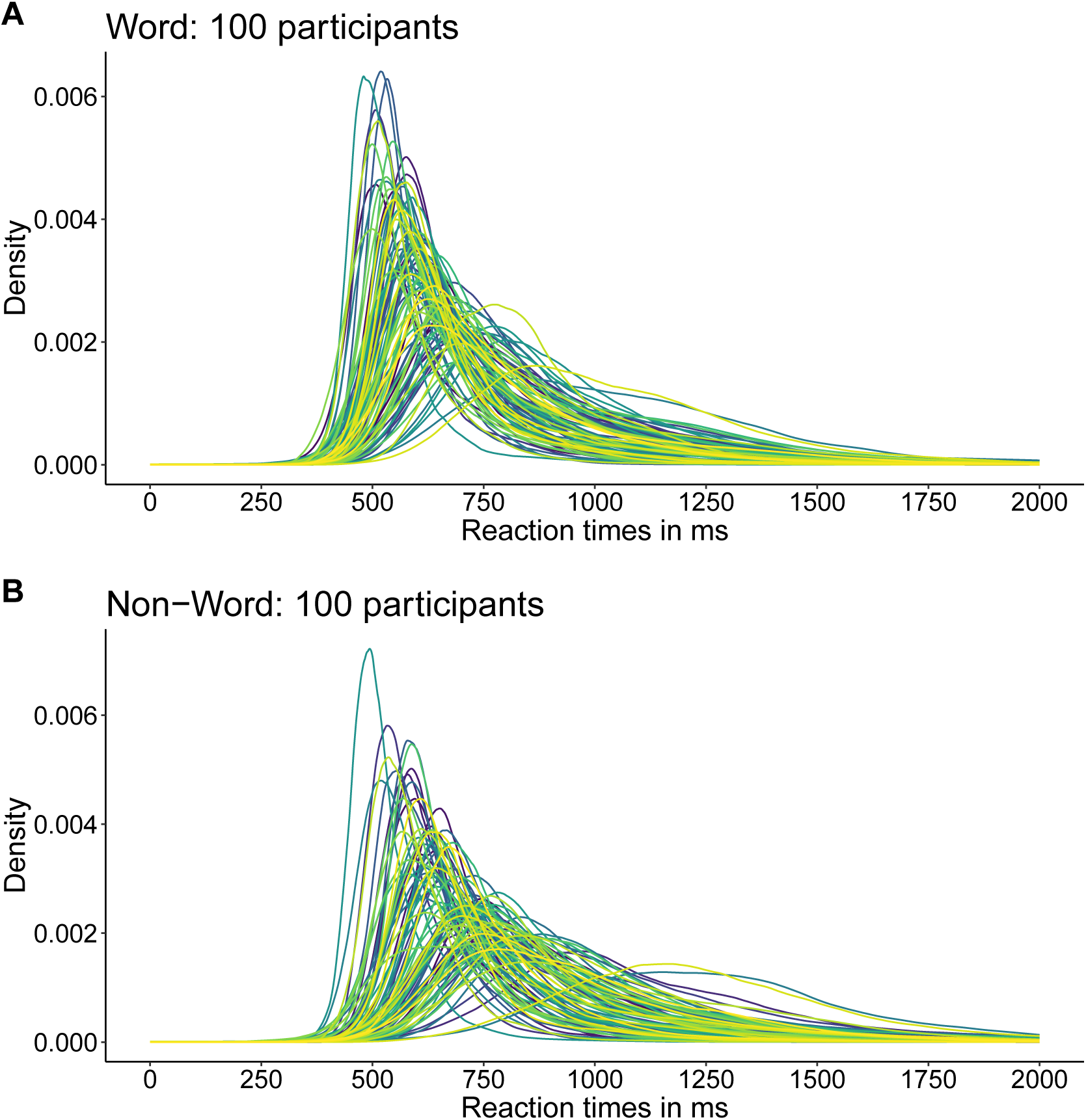
FLP dataset: reaction time distributions from 100 participants. Participants were randomly selected among 959. Distributions are shown for each participant (colour coded) in the Word (**A**) and Non-Word (**B**) conditions.

Among participants, the variability in the shapes of the distributions is particularly striking. The shapes of the distributions also differed between the Word and the Non-Word conditions. In particular, skewness tended to be larger in the Word than the Non-Word condition. Based on the standard parametric definition of skewness, that was the case in 80% of participants. If we use a non-parametric estimate instead (mean – median), it was the case in 70% of participants. This difference in skewness between conditions implies that the difference between medians will be biased in individual participants.

If we save the median response time for each participant and each condition, we get two distributions that display positive skewness (Figure 18A). The same applies to distributions of means (Figure 18B). The distributions of pairwise differences between the Non-Word and Word conditions is also positively skewed (Figure 18C). Notably, most participants have a positive difference: based on the median, 96.4% of participants are faster in the Word than the Non-Word condition; 94.8% for the mean.

**Figure 18.**
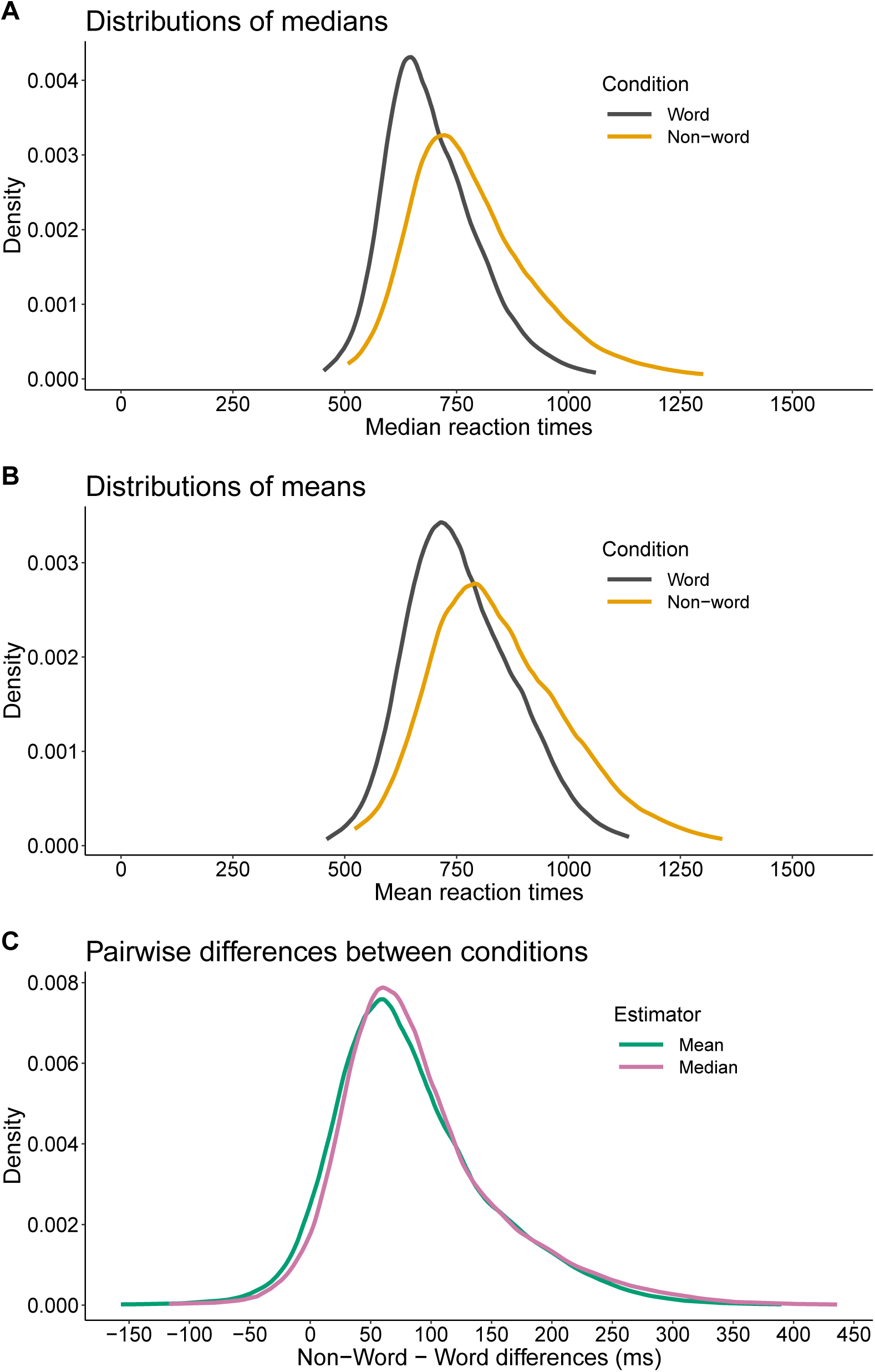
FLP dataset: group distributions. For every participant, the median (**A**) and the mean (**B**) were computed for the Word and Non-Word observations separately. Panel (**C.**) Distributions of pairwise differences between the Non-Word and Word conditions.

Hence, Figure 17 and Figure 18 demonstrates that we have to worry about skewness at 2 levels of analysis: in individual distributions and in group distributions. Here we explore estimation bias as a result of skewness and sample size in individual distributions, because it is the most similar to Miller’s 1988 simulations. Later we will consider false and true positives in a hierarchical situation. From what we’ve learnt so far, we can already make predictions: because skewness tended to be stronger in the Word than in the Non-Word condition, the bias of the median will be stronger in the former than the later for small sample sizes. That is, the median in the Word condition will tend to be more over-estimated than the median in the Non-Word condition. As a consequence, the difference between the median of the Non-Word condition (larger RT) and the median of the Word condition (smaller RT) will tend to be under-estimated. To check this prediction, we estimated bias in every participant using a simulation with 2,000 iterations (see details in notebook *flp bias sim*). We used the full sample of roughly 1,000 trials as the population, from which we computed population means and population medians. Because the Non-Word condition is the least skewed, we used it as the reference condition, which always had 200 trials. The Word condition had 10 to 200 trials, with 10 trial increments. In the simulation, single RT were sampled with replacements among the roughly 1,000 trials available per condition and participant, so that each iteration is equivalent to a fake experiment.

Let’s look at the results for the median. Figure 19A shows the bias of the difference between medians (Non-Word – Word), as a function of sample size in the Word condition. The Non-Word condition always had 200 trials. All participants are superimposed and shown as coloured traces. The average across participants is shown as a thicker black line.

**Figure 19.**
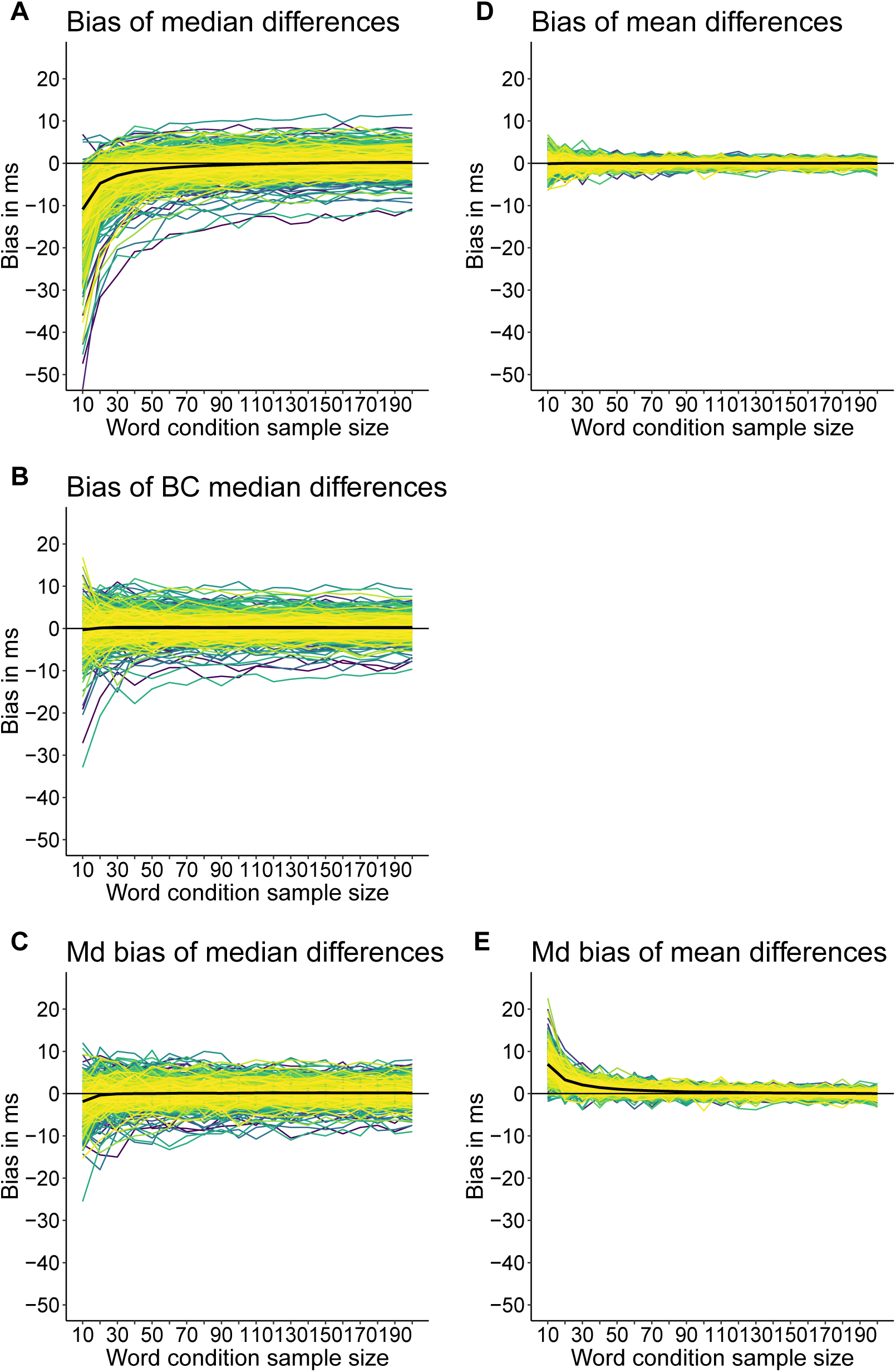
FLP dataset: bias estimation for the difference between the Non-Word and Word conditions. In each panel, thin coloured lines indicate results from individual participants and the thick black line indicates the mean across participants (group bias). The left column illustrates results for the median, the right column for the mean. BC = bias-corrected. Md bias = median bias.

As expected, bias tended to be negative with small sample sizes; that is, the difference between Non-Word and Word was underestimated because the median of the Word condition was overestimated. For the smallest sample size, the average bias was - 10.9 ms. That’s probably substantial enough to seriously distort estimation in some experiments. Also, variability is high, with a 80% highest density interval of [-17.1, −2.6] ms. Bias decreases rapidly with increasing sample size. For n=20, the average bias was −4.8 ms, for n=60 it was only −1 ms.

After bootstrap bias correction (with 200 bootstrap samples), the average bias dropped to roughly zero for all sample sizes (Figure 19B). Bias correction also reduced inter-participant variability.

As we saw in Figure 6, the sampling distribution of the median is skewed, so the standard measure of bias (taking the mean across simulation iterations) does not provide a good indication of the bias we can expect in a typical experiment. If instead of the mean, we compute the median bias, we get the results in Figure 19C. At the smallest sample size, the average bias is only −1.9 ms, and it drops to −0.3 for n=20. This result is consistent with the simulations reported above and confirms that in the typical experiment, the bias associated with the median is negligible.

What happens with the mean? The average bias of the mean is near zero for all sample sizes (Figure 19D). As we did for the median, we also considered the median bias of the mean (Figure 19E). For the smallest sample size, the average bias across participants is 6.9 ms. This positive bias can be explained from the results using the ex-Gaussian distributions: because of the larger skewness in the Word condition, the sampling distribution of the mean was more positively skewed for small samples in that condition compared to the Non-Word condition, with the bulk of the bias estimates being negative. That is, the mean tended to be more under-estimated in the Word condition, leading to larger Non-Word – Word differences in the typical experiment.

The results from the real RT distributions confirm our earlier simulations using ex-Gaussian distributions: for small sample sizes, the mean and the median differ in bias due to differences in sampling distributions and the bias of the median can be corrected using the bootstrap. Another striking difference between the mean and the median is the spread of bias values across participants, which is much larger for the median than the mean. This difference in bias variability does not reflect a difference in variability among participants for the two estimators of central tendency. Indeed, as we saw in Figure 18C, the distributions of differences between Non-Word and Word conditions are very similar for the mean and the median. Estimates of spread are also similar between difference distributions (median absolute deviation to the median (MAD): mean RT = 57 ms; median RT = 54 ms). This suggests that the inter-participant bias differences are due to the individual differences in shape distributions observed in Figure 17, to which the mean and the median are differently sensitive.

The larger inter-participant variability in bias for the median compared to the mean could also suggest that across participants, measurement precision would be lower for the median. We directly assessed measurement precision at the group level by performing a multi-level simulation. In this simulation, we asked, for instance, how often the group estimate was no more than 10 ms from the population value across many experiments (here 10,000 - see notebook *flp sim precision*). In each iteration (fake experiment) of the simulation, there were 200 trials per condition and participant, such that bias at the participant level was not an issue (a total of 400 trials for an RT experiment is perfectly reasonable and could be done in no more than 20-30 minutes per participant). For each participant and condition, the mean and the median were computed across the 200 random trials for each condition, and then the Non-Word - Word difference was saved. Group estimation of the difference was based on a random sample of 10 to 300 participants, with the group mean computed across participants’ differences between means and the group median computed across participants’ differences between medians. Population values were defined by first computing, for each condition, the mean and the median across all available trials for each participant, second by computing across all participants the mean and the median of the pairwise differences. Measurement precision was calculated as the proportion of experiments in which the group estimate was no more than *x* ms from the population value, with *x* varying from 5 to 40 ms.

Not surprisingly, the proportion of estimates close to the population value increases with the number of participants for the mean and the median (Figure 20). More interestingly, the relationship was non-linear, such that a larger gain in precision was achieved by increasing sample size for instance from 10 to 20 compared to from 90 to 100. The results also let us answer useful questions for planning experiments (see the black arrows in Figure 20A & B):

**Figure 20.**
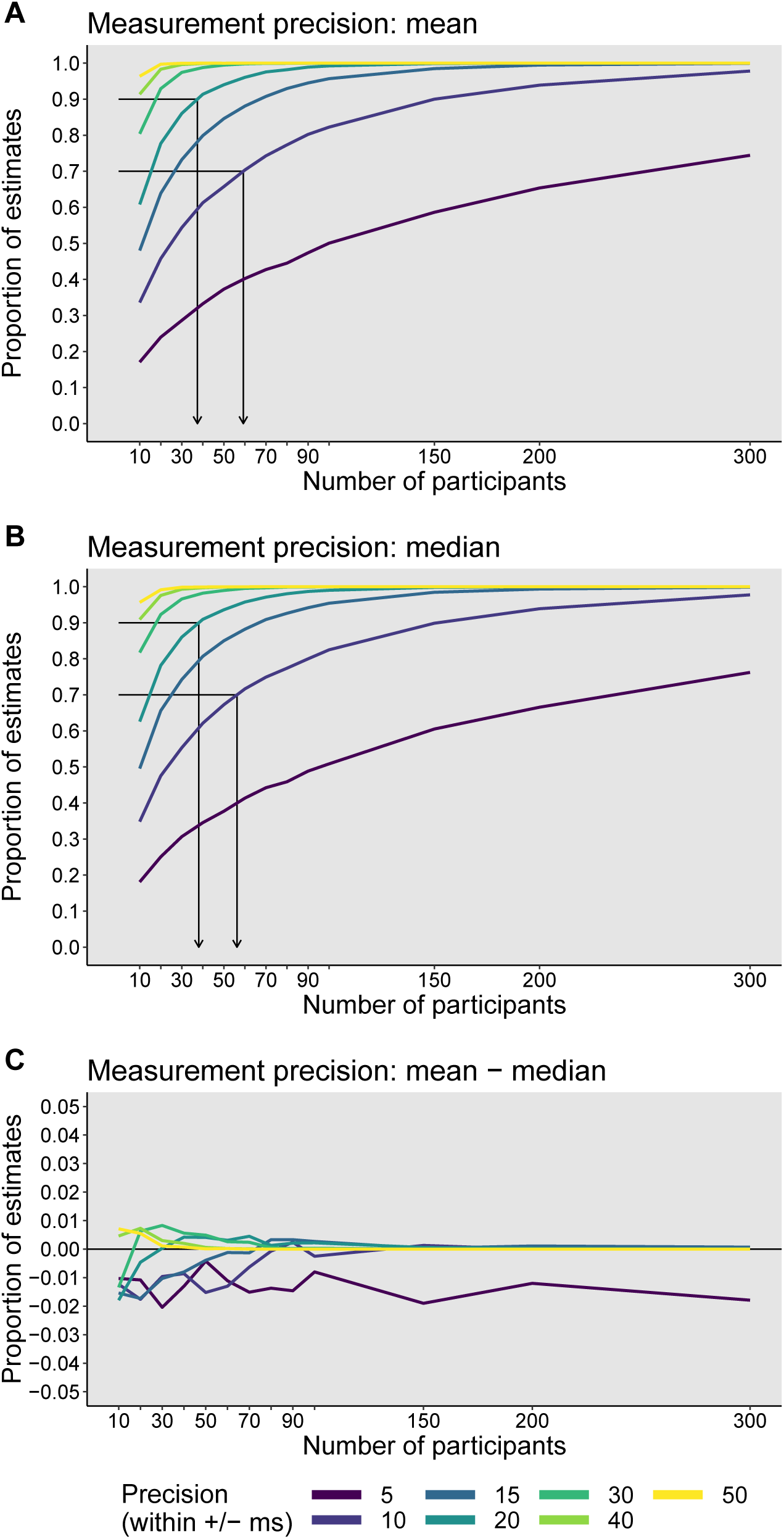
FLP dataset: group measurement precision for the difference between the Non-Word and Word conditions. Measurement precision was estimated by using a simulation with 10,000 iterations, 200 trials per condition and participant, and varying numbers of participants. Results are illustrated for the mean (**A**), the median (**B**), and the difference between the mean and the median (**C**).

- So that in 70% of experiments the group estimate is no more than 10 ms from the population value, we need to test at least 59 participants for the mean, 56 participants for the median.
- So that in 90% of experiments the group estimate is no more than 20 ms from the population value, we need to test at least 37 participants for the mean, 38 participants for the median.

Also, the mean and the median differed very little in measurement precision (Figure 20C), which suggests that at the group level, the mean does not provide any clear advantage over the median. Of course, different results could be obtained in different situations. For instance, the same simulations could be performed using different numbers of trials per participant and condition. Also, skewness can differ much more among conditions in certain tasks, such as in difficult visual search tasks (Palmer et al., 2011).

### False positives and true positives

Finally, we consider false positives and true positives using simulations with ex-Gaussian distributions. A limitation of previous ex-Gaussian simulations on false and true positives was the lack of inter-participant variability. To address this limitation, first we fit ex-Gaussian distributions to the Word and Non-Word conditions of each participant in the LFP dataset using maximum likelihood estimation (Massidda, 2013) - see notebook *flp exg parameters*. Then, we sample participants’ ex-Gaussian parameters with replacement, and use them to generate simulated trials from the Word and Non-Word conditions. Distributions of trials from each condition were summarised using the mean, the median and the deciles, as done previously. Distributions of individual pairwise differences were then tested against 0 using group tests on the mean of individual means and median of individual medians. For the deciles, we performed group tests on the mean, the median, the 10% and 20% trimmed means of each decile, with an Hochberg’s correction for multiple comparisons. For the deciles, the group statistics based on 20% trimmed means gave long-run proportions of false positives closer to the nominal level (see notebook *sim gp fp flp*). Using 20% trimmed means also gave the highest power in most situations, so we only report results for this group estimator (see full report in notebook *sim gp tp flp*).

For false positives, for each participant two samples were drawn using ex-Gaussian parameters from the Word condition, so that on average no effect was expected. Whether sample sizes were equal or not, group tests using means of means (mean), medians of medians (median) or a hierarchical shift function with 20% trimmed means (SF) gave proportions of false positives near the nominal level (Figure 21). With increasing numbers of participants the number of type I errors decreased slightly for the SF method, but remained well within the satisfactory range.

**Figure 21.**
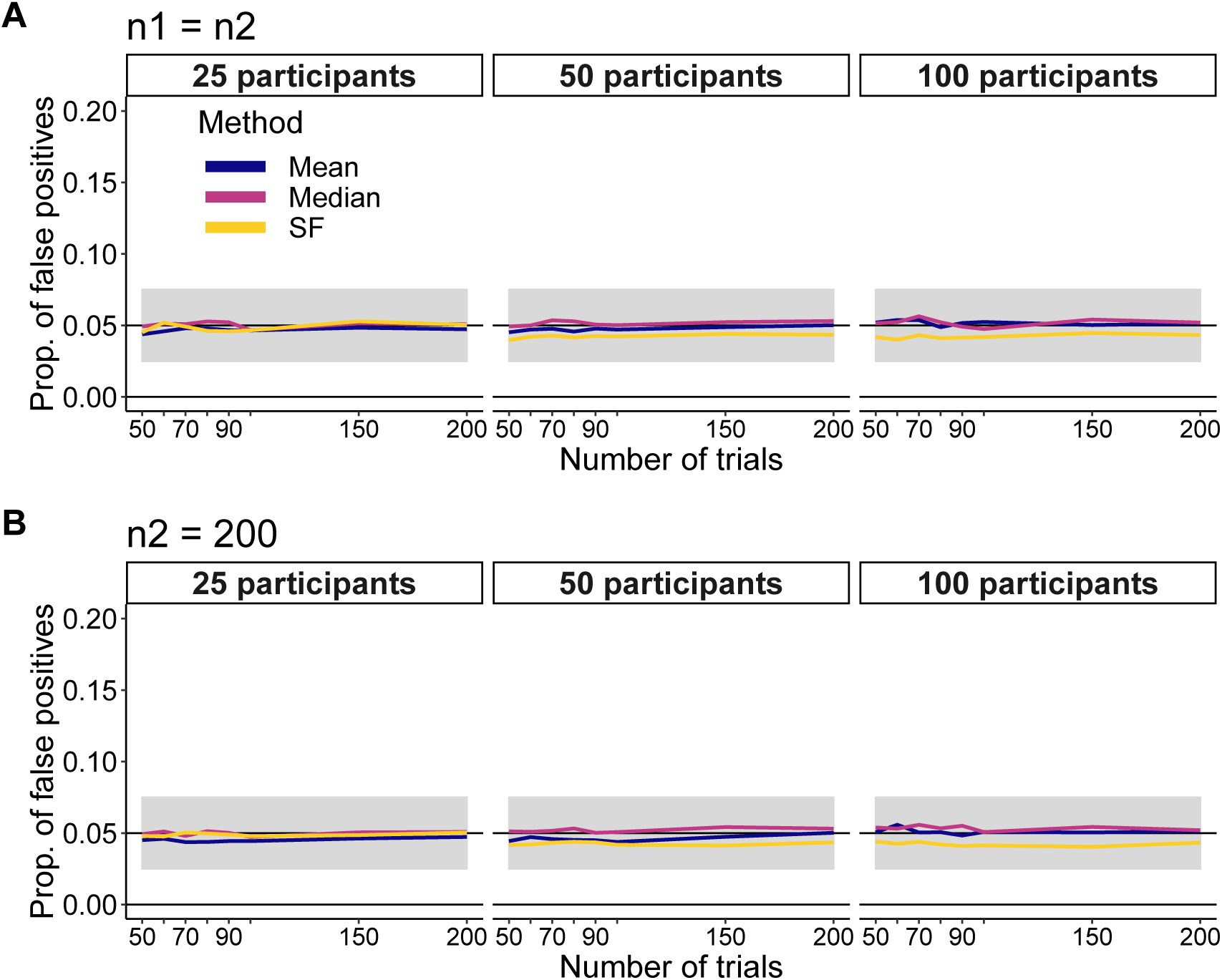
FLP dataset: false positive simulation. Results are shown for the cases in which each condition has the same number of trials (**A**) and different numbers of trials (**B**). Mean = means of means, Median = medians of medians, SF = hierarchical shift function with 20% trimmed means to assess group differences of deciles.

For true positives, for each participant we interpolated each ex-Gaussian parameter between the Word and Non-Word condition in 10 steps. For the simulated Word condition, we generated trials using the parameters from that condition; for the Non-Word condition, we used a combination of parameters from step 2 of the continuum. The results are shown in Figure 22. Across all conditions considered, the mean was associated with more power than the median. This can be explained by the relatively mild asymmetry of the distribution of pairwise differences for the mean and the median Figure 18: unlike the median, the mean is very sensitive to this asymmetry, but the asymmetry is not sufficient to inflate the mean’s standard error. In situations with larger asymmetry, it could become beneficial to use the median instead. Relative to the mean and the median, using a hierarchical shift function approach led to a large power increase across all trial and participant conditions.

**Figure 22.**
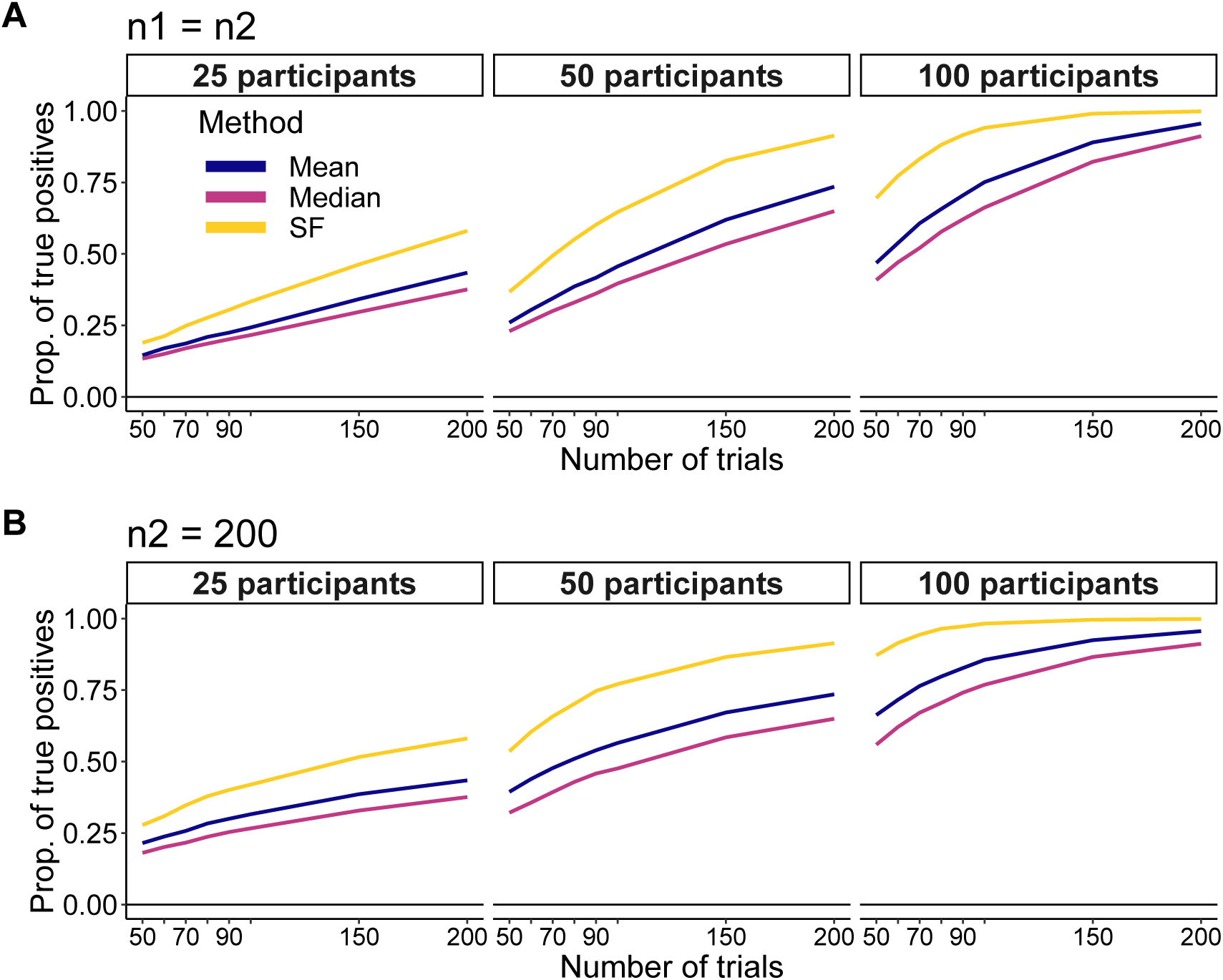
FLP dataset: true positive simulation. Results are shown for the cases in which each condition has the same number of trials (**A**) and different numbers of trials (**B**).

### A closer look

With the large effect sizes present in the FLP data, it wouldn’t matter whether we use the mean, the median, the hierarchical shift function technique or one of many other options: most would reject at low alpha levels. Over techniques considering only a measure of central tendency, the SF approach has the advantage of helping us understand the nature of the effects because it considers the shapes of the distributions. We can also get a better understanding of how distributions differ by plotting individual shift functions (Figure 23A). The 20% trimmed mean across participants is always positive and increases from early to late deciles: 59, 66, 72, 77, 82, 86, 89, 91, 89. Using Spearman’s correlation and an alpha of 0.05, 52.9% of participants were classified as showing a monotonic increase across deciles, and 14.9% showed a monotonic decrease. As illustrated in Figure 23B, at each decile most participants have a positive difference. The percentage of participants showing at every decile a positive difference was 83.2%; a negative difference 1.4%; the rest had deciles straddling the zero line. So a clear majority of participants showed stochastic dominance (Speckman, Rouder, Morey & Pratte, 2008). Whether there are really three groups with qualitatively different patterns of results across deciles could be addressed using recently proposed hierarchical models for instance (Haaf & Rouder, 2017). In sum, there is so much more to a data set than can be characterised by looking only at the mean or the median.

**Figure 23.**
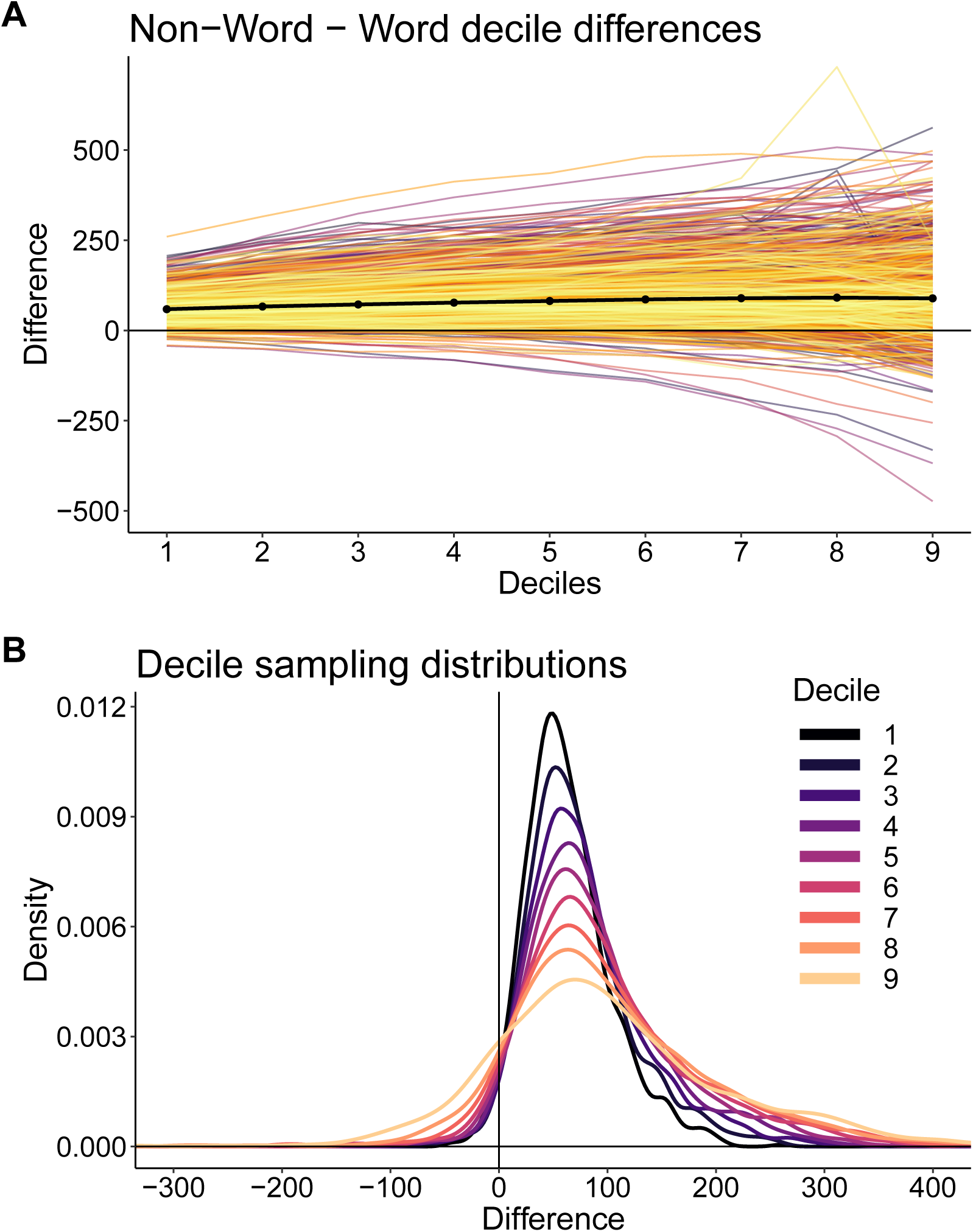
FLP dataset: illustrations of the deciles. (**A**) Hierarchical shift function. Each coloured line represent a participant. The thick black line shows the 20% trimmed mean across participants for each decile. (**B**) Sampling distributions of the deciles.

## Discussion

In this article, we reproduced the simulations from Miller (1988), extended them and applied a similar approach to a large dataset of reaction times (Ferrand et al., 2010). The two sets of analyses led to the same conclusion: the recommendation by Miller (1988) to not use the median when comparing distributions that differ in sample size was ill-advised, for several reasons. First, the bias can be strongly attenuated by using a percentile bootstrap bias correction. However, although the bootstrap bias correction appears to work well in the long-run, for a single experiment there is no guarantee it will provide an estimation closer to the truth. One possibility is to report results with and without bias correction. Second, the sample distributions of the mean and the median are positively skewed when sampling from positively skewed distributions such as RT distribution; as a result, computing the mean of the sample distributions to estimate bias can be misleading. If instead we consider the median of the sample distributions, we get a better indication of the expected bias in a typical experiment, and this median bias tends to be smaller for the sample median than the sample mean. Third, the mean and the median differ very little in the measurement precision they afford. Fourth, the mean is not robust to departures from normality and higher power is obtained with the median in many situations. Thus, there seems to be no rational for preferring the mean over the median as a measure of central tendency for skewed distributions. If the goal is to accurately estimate the central tendency of a RT distribution, while protecting against the influence of skewness and outliers, the median is far more efficient than the mean (Wilcox & Rousselet, 2018). Providing sample sizes are moderately large, bias is actually not a problem, and the typical bias is actually very small. So, given a choice between the mean and the median, the median appears to be a better option in some situations, at least for the theoretical and empirical distributions studied here, as which distributions best capture the shape of RT data is still debated (Campitelli, Macbeth, Ospina & Marmolejo-Ramos, 2017; Palmer et al., 2011). In other situations, the mean could be a better choice to detect differences, for instance in the presence of differences more strongly affecting late responses, or in the case of the moderately skewed distributions without clear outliers in the FLP data.

Clearly, no method dominates and researchers should use a range of tools to answer different questions about their data. Importantly, conclusions should be limited to the estimators used. For instance, when making inferences about the mean or the median, conclusions should be restrained to the mean or the median, not to the entire distributions. As we saw in our examples, distributions can differ in many ways and inferences on means or medians are limited because they ask specific, non-exhaustive questions about distribution differences.

We also need to keep in mind that, as we saw in the examples presented here, we have to make decisions about estimators at two levels of analysis: for participants and for the group. In our experience, these two choices are often ignored. For instance, in an extensive series of simulations, with 1886 citations as of January 10th 2019, Ratcliff (1993) demonstrated that when performing standard group ANOVAs, the median can lack power compared to other estimators. Ratcliff’s simulations involved ANOVAs on group means, in which for each participant and each condition, very few trials (7 to 12) were available. These small samples were summarised using several estimators, including the median. Based on the simulations, Ratcliff recommended data transformations or computing the mean after applying specific cut-offs to maximise power. However, these recommendations should be considered with caution because, first, the results could be very different with more realistic sample sizes, second, there is no rational for limiting group level analyses to mean differences. Like t-tests, standard ANOVAs on group means are not robust, and alternative techniques should be considered, involving trimmed means, medians and M-estimators (Field & Wilcox, 2017; Wilcox & Rousselet, 2018). More generally, standard procedures using the mean lack power, offer poor control over false positives, lead to inaccurate confidence intervals, and can have low estimation accuracy (Field & Wilcox, 2017; Wilcox & Rousselet, 2018; Davis-Stober, Dana & Rouder, 2018).

Ad-hoc data transformations suggested by Ratcliff (1993) are not ideal either, because they change the shapes of the distributions, which contain important information about the nature of the effects. Also, if transformations can successfully normalise distributions in some situations (Marmolejo-Ramos, Cousineau, Benites & Maehara, 2015), they sometimes fail to remove the skewness of the original distributions, and they do not effectively deal with outliers (Wilcox, 2017). Also, once data are transformed, inferences are made on the transformed data, not on the original ones, an important caveat that tends to be swept under the carpet when results are discussed. There is nevertheless a place for transformations chosen in a principled way to aid interpretation, for instance the log transformation for log distributed measurements.

Truncating distributions can also be detrimental, because it can introduce bias, especially when used in conjunction with the mean (Miller, 1991; Ulrich & Miller, 1994). Indeed, common outlier exclusion techniques lead to biased estimation of the mean (Miller, 1991). When applied to skewed distributions, removing any values more than 2 or 3 standard deviation from the mean affects slow responses more than fast ones. As a consequence, the sample mean tends to underestimate the population mean. And this bias increases with sample size because the outlier detection technique does not work for small sample sizes, which results from the lack of robustness of the mean and the standard deviation (Wilcox & Keselman, 2003). The bias also increases with skewness. Therefore, when comparing distributions that differ in sample size, or skewness, or both, differences can be masked or created, resulting in inaccurate quantification of effect sizes. Truncation using absolute thresholds (for instance by removing all RT *<* 300 ms and all RT *>* 1,200 ms and averaging the remaining values) also leads to potentially severe bias of the mean, median, standard deviation and skewness of RT distributions (Ulrich & Miller, 1994). The median is, however, much less affected by truncation bias than the mean. Also, the median is very resistant to the effect of outliers and can be used on its own without relying on dubious truncation methods. In fact, the median is a special type of trimmed mean, in which only one or two observations are used and the rest discarded (equivalent to 50% trimming on each side of the distribution). There are advantages in using 10% or 20% trimming in certain situations, and this can done in conjunction with the application of t-tests and ANOVAs for which the standard error terms are adjusted (Wilcox & Keselman, 2003; Wilcox, 2017).

Overall, there is no convincing evidence against using the median of RT distributions, if the goal is to use only one measure of location to summarise the entire distribution. Which to use, the mean or the median, depends on the empirical question and the type of data at hand. In many situations, we argue that neither should be used. Clearly, a better alternative is to not throw away all the information available in the raw distributions, by studying how entire distributions differ (Rousselet et al., 2017; Heathcote et al., 1991; Rouder & Province, Submitted; Baayen & Milin, 2010). This can be done for instance using the hierarchical shift function introduced in this article. Looking at multiple quantiles provides an effective way to boost power and, combined with detailed graphical representations, to understand how and by how much distributions differ. To be clear, we are not suggesting that the hierarchical shift function should be used in all situations. The choice of estimators and tests depends on the goal of the experimenter, and no method dominates. The particular implementations of the hierarchical shift function considered here is one of many potential candidates. It also has clear disadvantages over other methods. For instance, it does not include the shrinkage afforded by hierarchical models (Rouder & Province, Submitted). It is also blind to the underlying generative model, so it does not allow inferences more directly related to cognitive processes (Matzke et al., 2013; Voss, Nagler & Lerche, 2013). But this later point can be seen as an advantage, allowing flexible applications without having to specify a generative distribution. Also, very specific questions can be better answered by very specific tools. For instance, some methods have been proposed to quantify the minimal reaction times at which conditions differ (Rousselet et al., 2003; Reingold & Sheridan, 2018).

Whatever the approach, or conjunction of approaches chosen, we need to consider the shape of the distributions and outliers at both levels of analysis: in each participant and across participants. And these choices need to be justified. Using group means of individual means by default is not wise or justified. Finally, depending on the type of data and the goals of the experiment, an important question is how to spend our money: by investing in more trials or in more participants (Rouder & Haaf, 2018)? An answer can be obtained by running simulations, either data-driven using available large datasets or assuming generative distributions (for instance ex-Gaussian distributions for RT data). Simulations that take shape information into account are important to estimate bias and power. Assuming normality can have disastrous consequences (Wilcox & Rousselet, 2018). In many situations, simulations will reveal that much larger samples than commonly used are required to improve the precision of our estimations (Schönbrodt & Perugini, 2013; Peters & Crutzen, 2017; Rothman & Greenland, 2018).

## Contributions

- Contributed to conception and design: GAR, RRW
- Contributed to analysis and interpretation of data: GAR, RRW
- Drafted article: GAR
- Revised article: GAR, RRW
- Approved the submitted version for publication: GAR, RRW

## Acknowledgments

Our thanks to Fernando Marmolejo-Ramos and Jeff Miller for their feedback on the first preprint version of this article (Rousselet & Wilcox, 2018b). In particular Jeff Miller pointed out a mistake in the logic of one simulation and suggested we consider false positives in addition to bias.

## Data and code availability

All the figures in this article are licensed CC-BY 4.0 and can be reproduced using notebooks in the R programming language (R Core Team, 2018) and datasets available on *figshare* (Rousselet & Wilcox, 2018a): https://doi.org/10.6084/m9.figshare.6911924. The main R packages used to generate the data and make the figures are *ggplot2* (Wickham, 2016), *cowplot* (Wilke, 2017), *tibble* (Müller & Wickham, 2018), *tidyr* (Wickham & Henry, 2018), *retimes* (Massidda, 2013), *knitr* (Xie, 2018), *HDInterval* (Meredith & Kruschke, 2016), and the essential *beepr* (Bååth, 2018).

